# The mechanotransduction protein STOML3 is required for functional plasticity following peripheral nerve regeneration

**DOI:** 10.1101/2020.11.10.367748

**Authors:** Julia Haseleu, Jan Walcher, Gary R. Lewin

**Author notes:** The authors declare no competing financial interests.

## Abstract

Nerve regeneration is associated with plasticity of sensory neurons, so that even muscle afferents directed to skin form mechanosensitive receptive fields appropriate for the new target. STOML3 is an essential mechanotransduction component in many cutaneous mechanoreceptors. Here we asked whether STOML3 is required for functional and anatomical plasticity following peripheral nerve regeneration. We used a cross-anastomosis model adapted to the mouse in which the medial gastrocnemius nerve was redirected to innervate hairy skin previously occupied by the sural nerve. We recorded from muscle afferents innervating the skin and found that in wild-type mice their receptive properties were largely identical to normal skin mechanoreceptors. However, in mice lacking STOML3, muscle afferents largely failed to form functional mechanosensitive receptive fields, despite making anatomically appropriate endings in the skin. Our tracing experiments demonstrated that muscle afferents from both wild-type and *stoml3* mutant mice display remarkable anatomically plasticity, forming new somatotopically appropriate synaptic terminals in the region of the dorsal horn representing the sural nerve territory. The dramatic reduction in stimulus evoked activity from the cross-anastomosed gastrocnemius nerve in *stoml3* mutant mice did not prevent central anatomical plasticity. Our results have identified a molecular factor that is required for functional plasticity following peripheral nerve injury.

## Introduction

Sensory processing fundamentally relies on topographic mapping. Sensory inputs are spatially segregated and tiled to construct a somatotopic map within the dorsal horn that defines the location of the stimulus (Brown and Culberson, 1981; Shortland et al., 1989; Shortland and Woolf, 1993). After peripheral nerve transection and repair most sensory axons can reach their targets (Fawcett and Keynes, 1990; Tedeschi and Bradke, 2017), but do not regain their original topographical position in the skin (Burgess and Horch, 1973; Horch, 1979; Koerber et al., 1989; Lewin et al., 1994). Consequently, cutaneous receptive fields of dorsal horn neurons receiving input from regenerated fibres are initially large and diffuse (Lewin et al., 1994), but, after regeneration is complete, receptive fields shrink in an activity-dependent manner (Lewin et al., 1994). Since the pioneering work of Head and Rivers (Rivers and Head, 1908) it is known that regenerated sensory neurons form functional mechanosensory receptive fields (Burgess and Horch, 1973; Dykes and Terzis, 1979; Terzis and Dykes, 1980).

There has been much progress in investigating the molecular factors, both intrinsic and extrinsic, that drive regeneration of sensory axons (Chen et al., 2007; Tedeschi and Bradke, 2017; Mahar and Cavalli, 2018). For example HIF-1a was identified as a factors that can increase axon growth (Cho et al., 2015), but few studies have investigated how mechanosensitivity of regenerated axons is restored. Within hours after axotomy severed axons become mechanosensitive (Koschorke et al., 1994), a finding indicating that mechanosensitive ion channels are already present in regenerating sensory axons. Recently, the first molecular components of the sensory mechanotransduction apparatus have been identified in sensory neurons (Wetzel et al., 2007; Poole et al., 2014; Ranade et al., 2014). One of these molecules is the integral membrane protein stomatin-like protein-3 (STOML3) which is an essential component of the mechanotransduction complex in many mechanoreceptors (Wetzel et al., 2007, 2017) and works by dramatically increasing the sensitivity of mechanosensitive PIEZO2 channels (Poole et al., 2014). The *Piezo2* gene is required for normal touch sensation in humans (Chesler et al., 2016) and many mechanoreceptors and proprioceptors need this protein to respond to mechanical stimuli in mice (Ranade et al., 2014; Woo et al., 2015). Indeed, in both *stoml3* and *Piezo2* mutant mice around 40% of cutaneous myelinated sensory afferents completely lack mechanosensitivity (Wetzel et al., 2007, 2017; Ranade et al., 2014; Murthy et al., 2018). In this study we have addressed two related questions. First, does sensory mechanotransduction play a role in the functional recovery of regenerating axons when they reinnervate their original or a novel target after axonal injury? Second does activity driven by mechanical stimulation play a critical role in central plasticity after axonal injury?

Here we used a cross-anastomosis model in which the gastrocnemius nerve, a pure muscle nerve, is forced to regrow into the cutaneous sural nerve territory (McMahon and Gibson, 1987; McMahon and Wall, 1989). In this model muscle afferents are capable of functionally innervating foreign targets and gain receptive field properties appropriate for the new cutaneous target (Lewin and McMahon, 1991a, 1991b, 1993; Johnson et al., 1995). Muscle afferents that are forced to innervate the skin display substantial plasticity making new functional connections in the spinal cord appropriate for the new target (McMahon and Wall, 1989; Lewin and McMahon, 1993). Here we established this model in the mouse which allowed us to ask whether normal functional recovery after nerve regeneration requires the presence of the mechanotransduction molecule STOML3. Surprisingly, we found that STOML3 is required for most muscle afferents to make mechanosensitive endings in the skin. However, substantial central plasticity of the central terminals of these muscle afferents was still observed in the spinal cord. Our findings identify, for the first time, a molecular factor that is critical for the functional recovery of regenerating axons in the adult peripheral nervous system.

## Materials and Methods

Mice used in this study were *stoml3* mutant mice (Wetzel et al., 2007, 2017) which were bred over more than 10 generations onto a C57BL/6 background. The same C57BL/6 strain used for backcrossing was used as controls. All mice were housed and handled according to the German Animal Protection Law.

### Transganglionic tracing with cholera toxin subunit B conjugates

Four to five week old female and male mice were anesthetized by an intraperitoneal injection of ketamine (100 mg/kg) and xylazine (10 mg/kg). 0.2 μl of 1.5% cholera toxin subunit B conjugated with Alexa Fluor^®^ 594 (CTB; Thermo Fisher Scientific, Waltham, MA, USA) were injected subcutaneously into the second and third digit of the left and right hind paw, respectively, using a pulled glass pipette (5 μl PCR Pipets, Drummond Scientific Co, Broomall, PA, USA) attached to a Hamilton microliter syringe (Hamilton Bonaduz AG, Bonaduz, Switzerland). The glass capillary was inserted into the most distal interphalangeal crease an advanced under the skin towards the next proximal crease where the tracer was slowly injected. Five days post injection, allowing for transganglionic transport of the tracer, the mice were transcardially perfused with 0.1M phosphate buffered saline (PBS) and ice-cold 4% paraformaldehyde (PFA). Subsequently, tissues of interest (lumbar DRGs, spinal cord, and hind paw skin) were dissected out and postfixed overnight in 4% PFA at 4°C.

Mice subjected to cross-anastomosis surgeries were anesthetized 12 weeks post-surgery as described above. For intraneural CTB injections, the sural nerve was exposed in the popliteal fossa by an incision of the biceps femoris. To enable the insertion of a glass capillary, the nerve was freed from surrounding tissue and carefully lifted up by placing a small spatula under it. After puncturing its epineurium, the tip of a pulled glass capillary which was attached to a Hamilton microliter syringe was carefully inserted into the nerve and 2 μl of 1.5% CTB in 0.1 M PBS were slowly injected. Subsequently, the wound was washed out with 0.1 M PBS and closed using sterile sutures. The tracer was allowed to be transganglionically transported for five days after which the mice were transcardially perfused. Tissues of interest (spinal cord, peripheral nerves) were dissected out and postfixed overnight in 4% PFA at 4°C.

### Cross-anastomosis surgery

Four to five week old female and male mice were anesthetized by an intraperitoneal injection of ketamine (100 mg/kg) and xylazine (10 mg/kg). The sural nerve and the medial gastrocnemius nerve were exposed in the popliteal fossa by an incision of the biceps femoris, cut and cross-anastomosed as described before (McMahon and Gibson, 1987; Lewin and McMahon, 1991a). Briefly, the proximal stump of the sural nerve was joined to the distal stump of the medial gastrocnemius nerve and vice versa with an epineural suture stitch using swaged microsurgical sutures (11/0). On the contralateral site, the two nerves were either left intact, transected, or self-anastomosed. The wounds were washed out with 0.1 M PBS and closed in layers using sterile sutures. After 12 weeks, the mice were either sacrificed in order to perform skin-nerve preparation experiments or subjected to transganglionic tracing experiments.

### Tissue clearing

Routinely, fixed spinal cords were washed three times with 0.1 M PBS for 10 min each at room temperature (RT). Subsequently, they were immersed in ascending concentration series of 2,2’-thiodiethanol (TDE) for 24h each at RT (Staudt et al., 2007; Kloepper et al., 2010; Aoyagi et al., 2015; Costantini et al., 2015). The applied concentrations were 10%, 25%, 50%, and 97% TDE diluted with 0.1 M PBS. Alternatively, spinal cords were cleared using the optical clearing technique three-dimensional imaging of solvent-cleared organs (3DISCO) which is based on tetrahydrofuran (THF) and dibenzyl ether (DBE) or a combination of THF and TDE (Ertürk et al., 2011, 2012; Becker et al., 2013). Briefly, spinal cords were washed three times with 0.1 M PBS for 10 min each at RT. Subsequently, they were dehydrated and delipidated in 50%, 70%, and 80% THF diluted with ddH_2_O for 30 min each and three times in 100% THF for 30 min at RT. Then, the dehydrated tissues were either immersed in 100% DBE or in ascending concentration series of TDE.

Fixed whole-mount skin samples were washed three times with 0.1 M PBS for 10 min each at RT. Subsequently, the tissues were dehydrated and delipidated in 50%, 70%, and 80% THF diluted with ddH_2_O for 30 min each and three times in 100% THF for 30 min at RT. Finally, the dehydrated specimens were incubated in ascending concentration series of TDE for 12 hours each at RT.

During all incubation steps, the samples were kept on a vibrating table in the dark. For imaging, cleared spinal cords and whole-mount skin samples were mounted on glass slides in 97% TDE using press-to-seal silicone isolators (Electron Microscopy Sciences, Hatfield, PA, USA) with the dorsal surface facing up.

Fixed DRGs, peripheral nerves, and immunostained skin slices were cleared in ascending concentration series of TDE for a minimum of 120 min per concentration step and mounted on glass slides with coverslips.

### Immunostaining of thick skin slices

Dissected skin was postfixed in 4% PFA overnight. Subsequently, the skin was washed three times in PBS at RT for 10 min each and embedded in 3% low-melting agarose. Using a vibratome, the skin was cut into 100 μm thick transverse slices. Prior to antibody incubation, the skin slices were washed in blocking solution (5% normal serum, 0.1% Triton X-100 in 0.1 M PBS) at 4°C for 1 h. Skin slices were incubated with primary antibodies diluted in blocking solution at 4°C for 24 h. Next, the slices were washed three times in 0.1 M PBS for 10 min each at RT and incubated with secondary antibodies diluted in blocking solution at 4°C for 24 h. All incubation steps were performed under agitation and in the dark. After completed immunostaining, skin slices were optically cleared. Primary antibodies used were: chicken antiNF200 (Abcam, Cat# ab72996, RRID: AB_2149618) 1:2000, rabbit antiPGP9.5 (Dako, Cat# Z5116, RRID: AB_2622233) 1:500, rabbit antiS100 (Dako, Cat# Z0311, RRID: AB_10013383) 1:1000, rat antiCytokeratin8/18 (TROMA-I; DSHB, Cat# TROMA-1, RRID: AB_531826) 1:1000. Secondary antibodies (Invitrogen) were coupled to Alexa Fluor^®^ dyes (488, 647) and used at a dilution of 1:1000.

### Two-photon microscopy

Two-photon imaging was performed using a laser scanning microscope (LSM710 NLO; Carl Zeiss, Oberkochen, Germany) equipped with a tunable Ti:sapphire laser (Chameleon; Coherent, Santa Clara, CA, USA). Two channels were recorded sequentially to collect Alexa Fluor^®^ 594 fluorescence (excitation wavelength: 840 nm; emission range: 589-735 nm) and tissue autofluorescence (excitation wavelength: 780 nm; emission range: 504-608 nm). A 25× multi-immersion objective (0.8 numerical aperture) was used with water for uncleared sample imaging and with immersion oil for cleared sample imaging. Tiled stacks were taken through the spinal cord dorsal horn (for images shown in Figure 2 pixel size was 0.55 × 0.55 μm, z step size was 3 μm; for images shown in Figure 7 pixel size was 1 × 1 μm, z step size was 2 μm) and DRGs (pixel size: 1 × 1 μm, z step size: 4 μm).

### Confocal microscopy

Cleared whole-mount and sectioned skin samples were imaged using a laser scanning microscope (Zeiss LSM 710 NLO, Carl Zeiss, Oberkochen, Germany) equipped with a 10× objective (0.3 numerical aperture) and a 25× objective (0.8 numerical aperture). Fluorescence and transillumination images were acquired simultaneously.

### Electron microscopy

For electron microscopy, mice were transcardially perfused with 0.1 M PBS and ice-cold 4 % PFA. Nerves were dissected and postfixed in 4 % PFA/2.5 % glutaraldehyde in 0.1 M PBS for 3 days. Following treatment with 1% OsO_4_ in 0.1 M PBS for 2h at RT, the nerves were washed two times in 0.1 M PBS, dehydrated in a graded ethanol series and propylene oxide, and embedded in Poly/Bed^®^ 812 (Polysciences Europe GmbH, Hirschberg an der Bergstraβe, Germany). Semithin sections were stained with toluidine blue. Ultrathin sections (70 nm) were contrasted with uranyl acetate and lead citrate. Sections were examined with a Zeiss 910 electron microscope (Carl Zeiss, Oberkochen, Germany) and digital images were taken with a high-speed slow-scan CCD camera (Proscan, Lagerlechfeld, Germany) at an original magnification of ×1600. Three ultrathin sections were taken from three nerves and on each ultrathin section five images (16.83 μm × 12.91 μm) were taken. Myelinated axons were counted in these areas using ImageJ (Schneider et al., 2012). Axon counts were normalized to the whole nerve.

### Image processing

All images were processed using ImageJ (Schneider et al., 2012). Tiled stacks were stitched using either the imaging software ZEN 2010 (Carl Zeiss, Oberkochen, Germany) or the ImageJ plugin ‘Stitching 2D/3D’ (Preibisch et al., 2009). Subsequently, images were cropped to the same size and reduced to the same slice number. Next, background fluorescence was reduced by subtracting the autofluorescence channel from the CTB channel. Using stack histogram-based thresholding, the image stacks were binarised. The threshold was set as the mean grey value plus three times the standard deviation. Finally, single pixels were removed to reduce noise, e.g. hot pixels (Video 1).

In order to enable comparative analyses of spinal terminal fields of cutaneous myelinated afferents, the three-dimensional centres of mass of the voxel clouds representing CTB-labelled fibre terminals were determined using the ImageJ plugin ‘3D ImageJ Suite’ (Ollion et al., 2013). All images were aligned to the centre of mass of the voxel cloud, i.e. images were cropped to the same size and reduced to the same slice number around the respective centres of mass.

Summed dorso-ventral, rostro-caudal and/or medio-lateral projections of the binary image stacks were constructed to enable two-dimensional visualization of terminal fields (Video 1).

### Image analysis

Relative locations of spinal terminal field foci were determined with respect to the dorsal and medial grey/white matter border. The distance between the terminal field’s centre of mass and the dorsal as well as medial grey/white matter border was measured in summed rostro-caudal and dorso-ventral projections of binary image stacks, respectively.

Medio-lateral, rostro-caudal, and dorso-ventral spans of the terminal fields were measured in summed dorso-ventral (medio-lateral and rostro-caudal spans) and rostro-caudal (dorso-ventral span) projections of binary image stacks. Summed projections were thresholded with the threshold being set as the mean grey value plus one standard deviation. Subsequently, the dimensions of the bounding rectangles enclosing all pixels representing CTB-labelled fibre terminals were measured.

Areal densities (as voxels per area) of spinal terminal fields were calculated in summed dorso-ventral, medio-lateral, and rostro-caudal projections of binary image stacks. Using the ImageJ plugin ‘3D ImageJ Suite’, the total number of voxels representing CTB-labelled fibre terminals was determined in binary image stacks. Subsequently, the number of voxels was divided by the area (in μm^2^) that was occupied by positive pixels in summed dorso-ventral, medio-lateral, and rostro-caudal projections, respectively.

### *Ex vivo* skin nerve preparation studies

Ex vivo skin nerve preparations were performed as described before (Moshourab et al., 2013; Walcher et al., 2018). Briefly, mice were sacrificed and the hair on the left hind limb was removed. The sural nerve (intact or regenerated) or the rerouted medial gastrocnemius nerve, respectively, was exposed in the popliteal fossa and dissected free along the lower leg. Subsequently, the skin was carefully removed from the musculoskeletal and connective tissue of the paw. The skin-nerve preparation was placed in an organ bath filled with oxygenated 32°C warm synthetic interstitial fluid (SIF; NaCl, 123 mM; KCl, 3.5 mM; MgSO_4_ mM, 0.7; NaH_2_PO_4_ mM, 1.7; CaCl_2_, 2.0 mM; sodium gluconate, 9.5 mM; glucose, 5.5 mM; sucrose, 7.5 mM; and HEPES, 10 mM at a pH of 7.4). Using insect needles, the skin was mounted in the organ bath with its epidermis facing the bottom of the chamber, exposing the dermis to the solution. The nerve was pulled through a hole into the adjacent recording chamber which was filled with mineral oil. Finally, using fine forceps the nerve was desheathed by removing its epineurium and small filaments were teased of the nerve. Throughout the whole experiment the skin was superfused with oxygenated SIF at a flow rate of 15 ml/min.

Teased filaments were attached to a recording electrode. The receptive fields of individual units were identified by manually probing the skin with a glass rod. Subsequently, the units were classified as RAMs, SAMs, D-hair receptors, and AMs, respectively, based on their conduction velocity (CV), spike pattern, and sensitivity. For immediate visual identification of single units, whole action potential waveforms were resolved on an oscilloscope. Data was acquired using a PowerLab 4/30 system (ADInstruments Ltd, Oxford, UK) which was controlled with the software LabChart 7.1 (ADInstruments Ltd, Oxford, UK).

The CVs of single fibres were determined by evoking a local action potential with a platinum iridium electrode (1 MΩ; World Precision Instruments Germany GmbH, Berlin, Germany). The electrical impulse was conducted nearly instantaneously through the solution whereas the triggered action potential conducted by the fibre was delayed depending on the fibre type and the distance of the fibre’s receptive field from the electrode. Hence, the distance between the receptive field of a unit to the electrode was measured and the CV was calculated as distance divided by time delay. Routinely, fibres with CVs above 10 m/s were classified as Aβ-fibres and those with CVs between 1.5 m/s and 10 m/s were classified as Aδ-fibres.

Mechanically sensitive receptors were stimulated using either a piezo actuator (Physik Instrumente GmbH & Co KG, Karlsruhe Germany) delivering dynamic and vibratory stimuli or a nanomotor (Kleindieck Nanotechnik GmbH, Reutlingen, Germany) enabling static stimulations. Both, the piezo actuator and the nanomotor were connected to a force sensor and mounted on a manual micromanipulator. Based on their response properties to various ramp-and-hold stimuli, Aβ- and Aδ-fibres were further classified as innervating RAMs or SAMs and D-hair receptors or AMs, respectively. Using the piezo actuator, dynamic mechanical stimuli were delivered in the form of ramp-and-hold stimuli with constant force (approximately 40 mN) but ramp phases of different velocities (0.075 mm/s, 0.15 mm/s, 0.45 mm/s, and 1.5 mm/s). Spikes elicited during the dynamic phase of the stimulus were analysed. Furthermore, sinusoidal vibration stimuli (25 Hz and 50 Hz) increasing in amplitude were given to determine the fibre’s mechanical threshold as the minimal force needed to evoke an action potential. Static mechanical stimuli were delivered using the nanomotor which was controlled by the NanoControl 4.0 software (Kleindieck Nanotechnik GmbH, Reutlingen, Germany). Ramp-and-hold stimuli with a constant ramp (1.5 – 2 mN/ms) but varying amplitudes were applied. Spikes evoked during the static phase of the stimulus were analysed.

For the electrical search protocol, a microelectrode (0.5-1 MΩ) was maneuvered gently to contact the epineurium of the nerve and electrical stimulations at 1 s intervals with square pulses of 50-500 ms duration were delivered. Electrically identified units were traced to their receptive fields. Subsequently, mechanical sensitivity of single units was tested by mechanical stimulation of their receptive field with a glass rod; units not responding to mechanical probing were designated as mechano-inensitive. Based on the CV, these units were categorized as mechano-insensitive Aβ- or Aδ-fibres.

### Statistical analysis

All statistical analyses were performed using the statistical software Prism 6 (GraphPad Software Inc, La Jolla, CA, USA). Depending on the experimental design, data sets were analysed using a two-tailed unpaired t-test (with Welch’s correction for data sets with unequal variances), one-way analysis of variance (ANOVA; with Tukey’s multiple comparison test), two-way repeated measures ANOVA (with Bonferroni post hoc test), or two-sided Fisher’s exact tests. In case where two-way repeated measures ANOVA were performed p-values for interaction effects are stated. Data sets were considered significantly different for p-values lower than 0.05. In figures, p-values are represented using the asterisk rating system where p < 0.05 is indicated by one asterisk (*), p < 0.01 by two asterisks (**), and p < 0.001 by three asterisks (***). All data are presented as mean ± standard error of the mean (SEM). Replicates are biological replicates.

## Results

### Structural and functional plasticity of primary sensory neurons

We investigated how mechanosensory silence affects structural and functional plasticity following nerve injury. We adapted a cross-anastomosis model to the mouse in which the medial gastrocnemius nerve, a pure muscle nerve, is cross-anastomosed to the cut cutaneous sural nerve which innervates the lateral hind paw and ankle (McMahon and Wall, 1989; Lewin and McMahon, 1991a, 1991b) (Figure 1a,b). Thus, muscle afferents of the medial gastrocnemius nerve are forced to regrow into the skin territory of the sural nerve and sural nerve sensory fibres are redirected to the medial gastrocnemius muscle via the distal cut end of the muscle nerve. In the rat and cat it was shown that muscle and cutaneous afferents redirected towards inappropriate targets, i.e. muscle afferents to skin and vice versa, gain neurochemical and physiological properties appropriate for their new target (McMahon and Gibson, 1987; McMahon and Wall, 1989; Lewin and McMahon, 1991a, 1991b; Johnson et al., 1995). We first performed cross-anastomosis surgeries in wild-type mice to investigate the capacity of muscle afferents to functionally innervate the skin.

**Figure 1.**
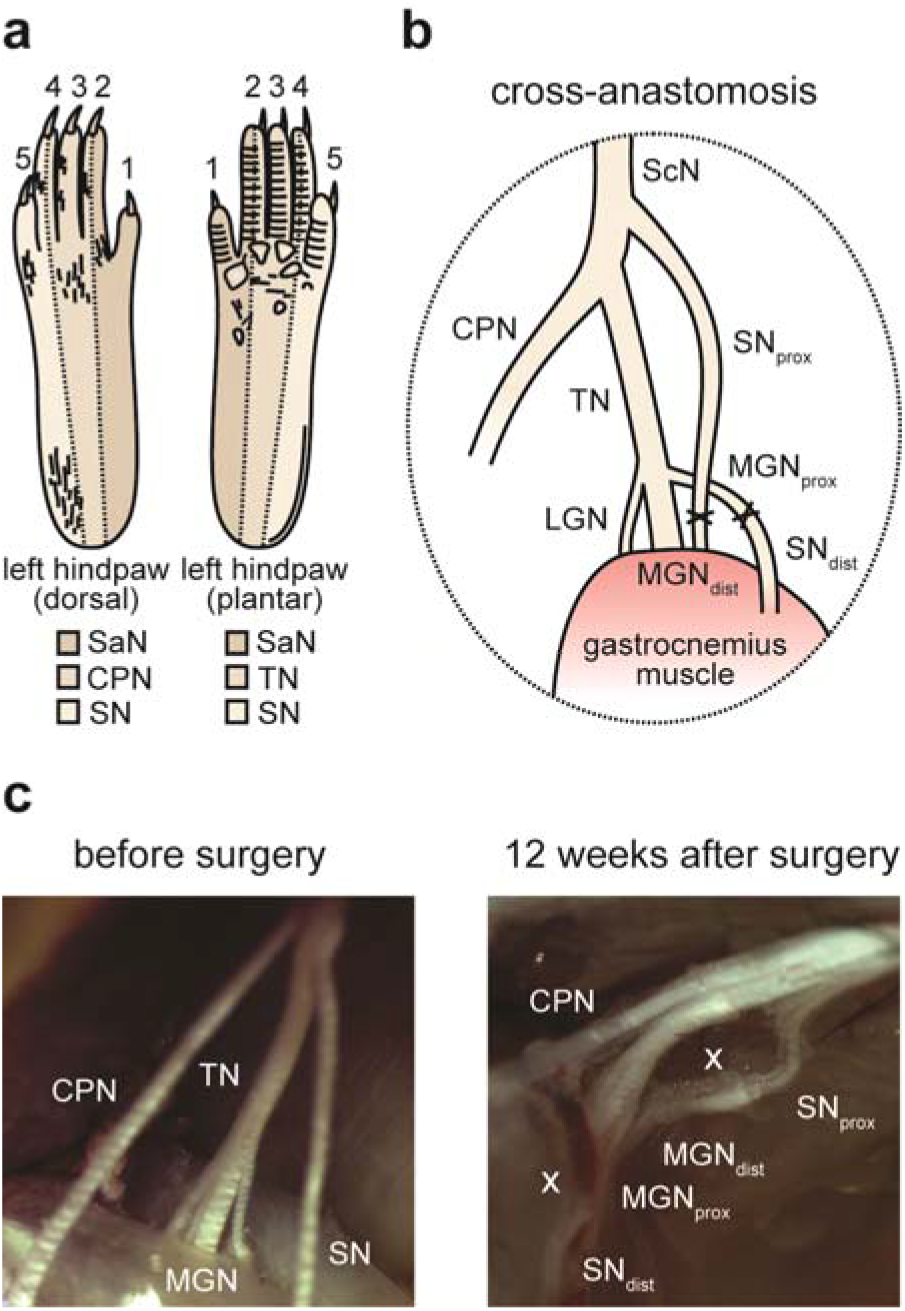
Cross-anastomosis of the sural and gastrocnemius nerve in the popliteal fossa. **(a)** Schematic representation of the innervation territories of four peripheral nerves, i.e. saphenous nerve (SaN), common peroneal nerve (CPN), tibial nerve (TN), and sural nerve (SN) in the left hind paw. **(b)** In the cross-anastomosis model, the sural nerve (SN) and the medial gastrocnemius nerve (MGN) are cross-anastomosed. **(c)** Stereomicroscopic images of the peripheral nerves innervating the hind paw in the popliteal fossa before and 12 weeks after cross-anastomosis surgery. The cross-anastomosis sites are marked with an ‘X’. Abbreviations: SaN, saphenous nerve; CPN, common peroneal nerve; SN, sural nerve; TN, tibial nerve; ScN, sciatic nerve; LGN, lateral gastrocnemius nerve; MGN, medial gastrocnemius nerve.

Cross-anastomosis surgeries were performed on four-week old wild-type mice. As a control, the sural nerve was either left intact or self-anastomosed. During terminal experiments (12 weeks post-surgery), the cross-anastomosed nerves were examined to exclude inappropriate nerve regeneration. In all cases, the cross-anastomosed nerves showed intact epineural sheaths and were clearly separable from each other as well as from other peripheral nerves within the popliteal fossa (Figure 1c).

We made extracellularly recordings from single intact and regenerated fibres in wild-type mice using the *ex vivo* skin nerve preparation adapted to the sural nerve territory. Consistent with previous studies in the cat and rat (Lewin and McMahon, 1991a; Johnson et al., 1995), single units with response properties characteristic of rapidly-adapting and slowly-adapting mechanoreceptors (RAMs and SAMs) (Figure 4a) as well as A-mechanonociceptors (AMs) (Figure 2b) were found in all three preparations (‘intact’, ‘self’, and ‘cross’). Functional D-hair receptors were only found in preparations of the intact and regenerated sural nerve, but not in preparations where muscle afferents were redirected towards the skin (Figure 2b). In the cross-anastomosed gastrocnemius nerve one Aβ-fibre (1/18) was found that only responded to manually delivered rapid and vigorous tapping of the receptive field, a rapid change in force beyond what could be delivered by the electromechanical stimulator (> 1.5 mm/s) (Figure 2a). This type of mechanically insensitive fibre type (tap-unit) has been noted rarely in wild-type nerves, but was found to be more frequent in *stoml3* mutant mice (Wetzel et al., 2007; Moshourab et al., 2013). Table 1 provides an overview of the number of characterized units in the three preparations and their conduction velocities. As expected from the results of previous studies (Horch and Lisney, 1981; Lewin and McMahon, 1991a; Johnson et al., 1995), the conduction velocities of Aβ-fibres were significantly slower in the regenerated nerves (both self- and cross-anastomosed nerves) when compared to those recorded from the intact nerve (Table 1). In contrast, the conduction velocities of Aδ-fibres were not significantly different between any of the three experimental groups (Table 1).

**Table 1.**
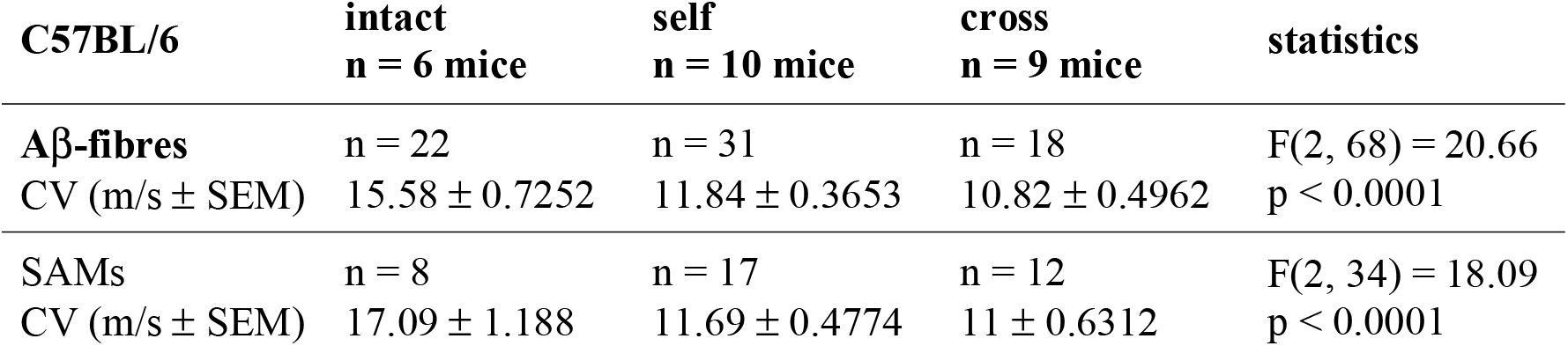
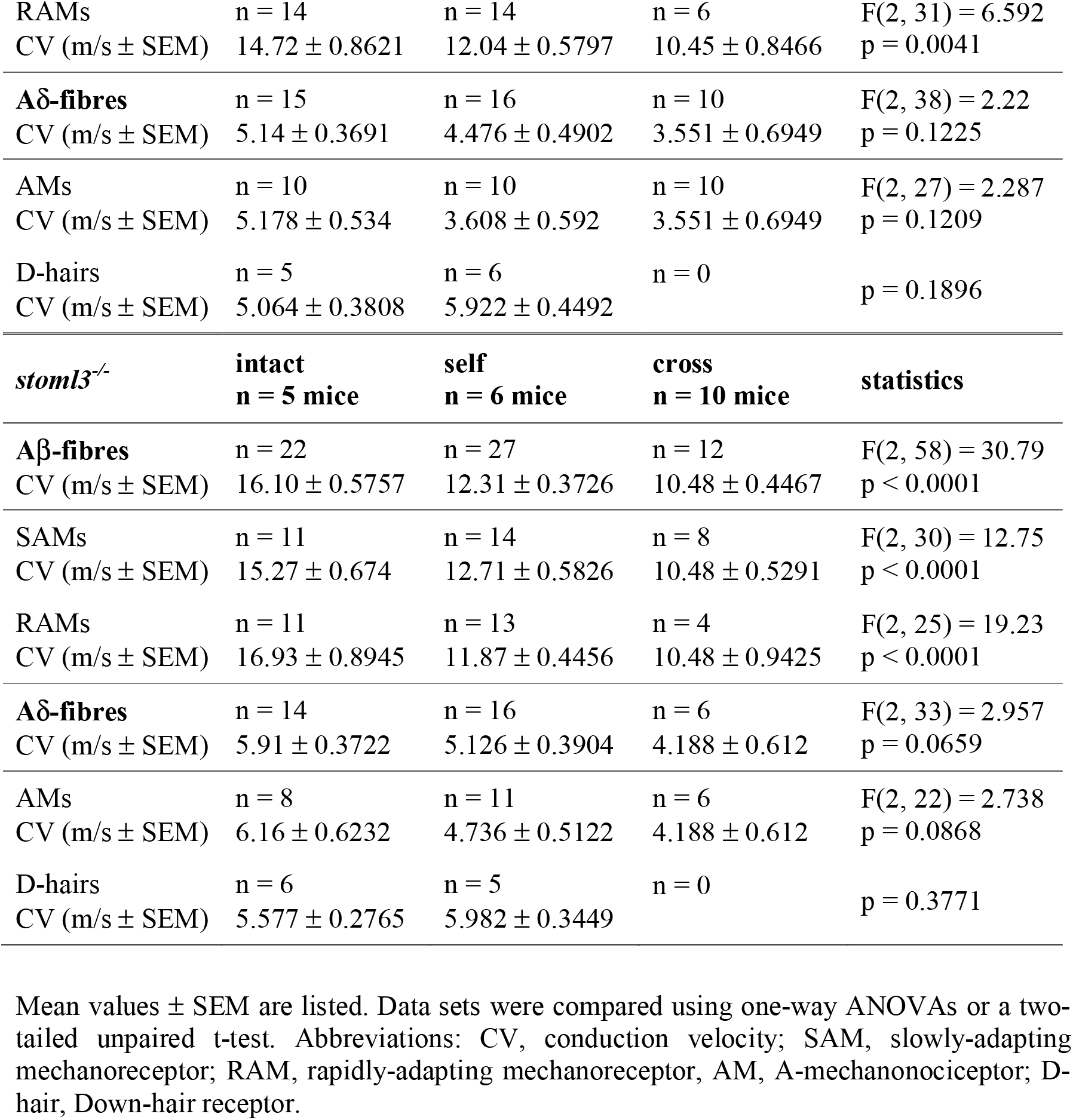
Conduction velocities of intact cutaneous, regenerated cutaneous, and redirected muscle afferents innervating hind paw skin in C57BL/6 and *stoml3* mutant mice.

| C57BL/6 | intact<br>n = 6 mice | self<br>n = 10 mice | cross<br>n = 9 mice | statistics |
| --- | --- | --- | --- | --- |
| **Aβ-fibres** | n = 22 | n = 31 | n = 18 | $F(2, 68) = 20.66$ |
| CV (m/s ± SEM) | $15.58 \pm 0.7252$ | $11.84 \pm 0.3653$ | $10.82 \pm 0.4962$ | $p < 0.0001$ |
| **SAMs** | n = 8 | n = 17 | n = 12 | $F(2, 34) = 18.09$ |
| CV (m/s ± SEM) | $17.09 \pm 1.188$ | $11.69 \pm 0.4774$ | $11 \pm 0.6312$ | $p < 0.0001$ |
| RAMs | n = 14 | n = 14 | n = 6 | F(2, 31) = 6.592 |
| CV (m/s ± SEM) | 14.72 ± 0.8621 | 12.04 ± 0.5797 | 10.45 ± 0.8466 | p = 0.0041 |
| **Aδ-fibres** | n = 15 | n = 16 | n = 10 | F(2, 38) = 2.22 |
| CV (m/s ± SEM) | 5.14 ± 0.3691 | 4.476 ± 0.4902 | 3.551 ± 0.6949 | p = 0.1225 |
| AMs | n = 10 | n = 10 | n = 10 | F(2, 27) = 2.287 |
| CV (m/s ± SEM) | 5.178 ± 0.534 | 3.608 ± 0.592 | 3.551 ± 0.6949 | p = 0.1209 |
| D-hairs | n = 5 | n = 6 | n = 0 | p = 0.1896 |
| CV (m/s ± SEM) | 5.064 ± 0.3808 | 5.922 ± 0.4492 | | |
| *stoml3*<sup>-/-</sup> | **intact**<br>**n = 5 mice** | **self**<br>**n = 6 mice** | **cross**<br>**n = 10 mice** | **statistics** |
| **Aβ-fibres** | n = 22 | n = 27 | n = 12 | F(2, 58) = 30.79 |
| CV (m/s ± SEM) | 16.10 ± 0.5757 | 12.31 ± 0.3726 | 10.48 ± 0.4467 | p < 0.0001 |
| SAMs | n = 11 | n = 14 | n = 8 | F(2, 30) = 12.75 |
| CV (m/s ± SEM) | 15.27 ± 0.674 | 12.71 ± 0.5826 | 10.48 ± 0.5291 | p < 0.0001 |
| RAMs | n = 11 | n = 13 | n = 4 | F(2, 25) = 19.23 |
| CV (m/s ± SEM) | 16.93 ± 0.8945 | 11.87 ± 0.4456 | 10.48 ± 0.9425 | p < 0.0001 |
| **Aδ-fibres** | n = 14 | n = 16 | n = 6 | F(2, 33) = 2.957 |
| CV (m/s ± SEM) | 5.91 ± 0.3722 | 5.126 ± 0.3904 | 4.188 ± 0.612 | p = 0.0659 |
| AMs | n = 8 | n = 11 | n = 6 | F(2, 22) = 2.738 |
| CV (m/s ± SEM) | 6.16 ± 0.6232 | 4.736 ± 0.5122 | 4.188 ± 0.612 | p = 0.0868 |
| D-hairs | n = 6 | n = 5 | n = 0 | p = 0.3771 |
| CV (m/s ± SEM) | 5.577 ± 0.2765 | 5.982 ± 0.3449 | | |

We next assessed the stimulus-response functions of muscle afferents innervating the skin compared to those of regenerated and intact cutaneous afferents. Normal RAMs are primarily tuned to stimulus velocity (Walcher et al., 2018), and so we used a series of increasing velocity stimuli (ramp-and-hold stimuli with probe velocities of 0.075 mm/s, 0.15 mm/s, 0.45 mm/s, and 1.5 mm/s) at a constant displacement of 96 μm to probe mechanoreceptor sensitivity. In addition, we used a sinusoidal vibration stimulus (50 Hz) applied with increasing amplitude to determine the minimal mechanical threshold to activate mechanoreceptors. No significant differences in the stimulus-response properties of RAMs were observed across the three experimental groups (Figure 2c, example traces are shown); spike frequencies of RAMs in response to moving stimuli were essentially identical across the groups (‘intact’: n = 14 fibres; ‘self’: n = 14 fibres; ‘cross’: n = 6 fibres; two-way repeated measures ANOVA: F(6, 93) = 0.9800, p = 0.443; Figure 4d) as were mechanical thresholds (‘intact’: 13.9 ± 2.4 mN, n = 10 fibres; ‘self’: 7.1 ± 1.3 mN, n = 14 fibres; ‘cross’: 10.6 ± 5.2 mN, n = 5 fibres; one-way ANOVA: F(2, 26) = 2.543, p = 0.098; Figure 2e). The response properties of SAMs were examined using a series of increasing displacement stimuli with a constant ramp velocity. The displacements ranged from 15 mN to 250 mN and lasted 2 s during the hold phase. Spike frequencies (in Hz) were calculated by counting the number of spikes occurring during the hold phase of the stimulus. Mechanical thresholds (in mN) were assessed by measuring the minimal force needed to evoke an action potential, i.e. by measuring the force at which the first action potential occurred during the dynamic phase of the stimulus. In all three preparations (‘intact’, ‘self’, and ‘cross’), characteristic SAM responses were recorded (Figure 2f, example traces are shown). However, the stimulus-response functions of SAMs in cross-anastomosed gastrocnemius nerve preparations were significantly larger in the suprathreshold range compared to intact and self-anastomosed sural nerve preparations (‘intact’: n = 8 fibres; ‘self’: n = 17 fibres; ‘cross’: n = 12 fibres; two-way repeated measures ANOVA: F(6, 102) = 4.700, p < 0.001; Bonferroni post hoc test; Figure 2g). Super-sensitive SAM responses in the cross-anastomosed nerve might reflect the intrinsic properties of former muscle spindle afferents that can sustain extremely high firing rates. No significant differences in the mechanical thresholds of SAMs were observed between the three preparations (‘intact’: 7.5 ± 1.4 mN, n = 8 fibres; ‘self’: 7.7 ± 1.6 mN, n = 17 fibres; ‘cross’: 5.5 ± 1.1 mN, n = 12 fibres; one-way ANOVA: F(2, 34) = 0.6402, p = 0.533; Figure 2h).

No sensory fibres with physiological attributes of D-hair receptors were found in the cross-anastomosed gastrocnemius nerve. However, the stimulus-response properties of D-hair receptors found in intact and self-anastomosed sural nerves were not different (Figure 2i, example traces are shown). Both spike frequencies (‘intact’: n = 5 fibres; ‘self’: n = 6 fibres; two-way repeated measures ANOVA: F(3,27) = 1.124, p = 0.357; Figure 2, Figure 2-1, Figure 1a) and mechanical thresholds (‘intact’: 0.3 ± 0.1 mN, n = 5 fibres; ‘self’: 0.5 ± 0.2 mN, n = 5 fibres; two-tailed unpaired t-test: t(8) = 1.302, p = 0.229; Figure 2, Figure 2-1b) were essentially identical between intact and self-anastomosed sural nerves, indicating that D-hair receptors easily regain their functional properties following nerve lesion. Characteristic AM responses were found in all three preparations (Figure 2j, examples traces are shown). The response properties of AMs including spike frequencies (‘intact’: n = 10 fibres; ‘self’: n = 10 fibres; ‘cross’: n = 10 fibres; two-way repeated measures ANOVA: F(6,81) = 1.325, p = 0.255; Figure 2, Figure 2-1) and mechanical thresholds (‘intact’: 76.2 ± 11.3 mN, n = 10 fibres; ‘self’: 65.8 ± 11.8 mN, n = 10 fibres; ‘cross’: 57.3 ± 16.7 mN, n = 10 fibres; one-way ANOVA: F(2,27) = 0.4928, p = 0.616; Figure 2, Figure 2-1c,d) were not significantly different from each other.

**Figure 2.**
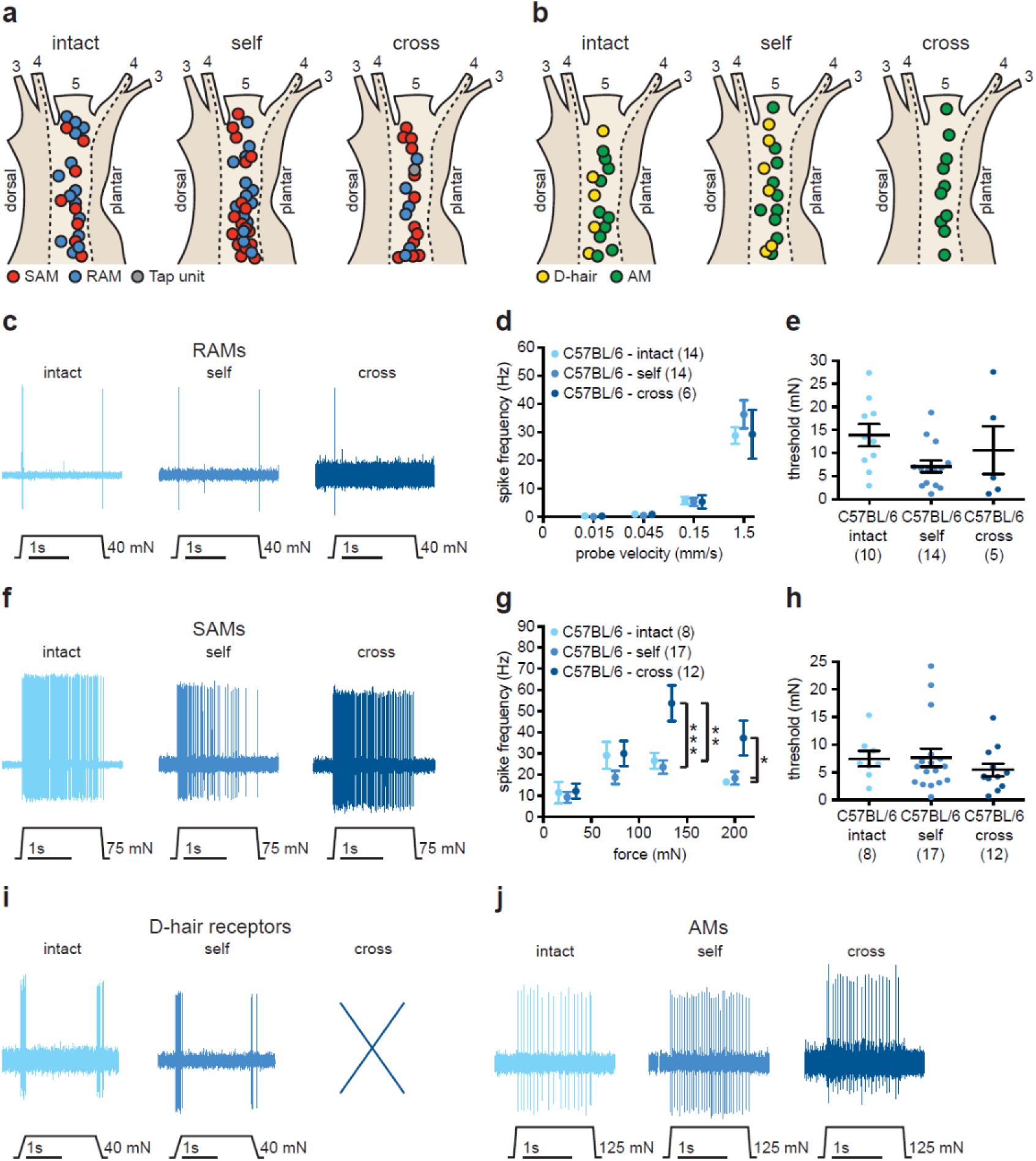
Response properties of muscle afferents newly innervating the skin compared to intact and regenerated cutaneous afferents in C57BL/6 mice. **(a,b)** Receptive field locations of intact cutaneous, regenerated cutaneous, and redirected muscle **(a)** Aβ-fibres (SAMs, red; RAMs, blue; tap-units, grey) and **(b)** Aδ-fibres (D-hair receptors, yellow; AMs, green). **(c)** Example traces of RAM responses to a ramp-and-hold stimulus with a probe velocity of 1.5 mm/s. **(d)** Spike frequencies of RAMs in response to ramp-and-hold stimuli with increasing ramp velocities. Mean values ± SEM. **(e)** Mechanical thresholds of RAMs measured in response to a sinusoidal vibration stimulus (50 Hz). Individual data points and mean values ± SEM are shown. **(f)** Example traces of SAM responses to a ramp-and-hold stimulus with an indentation force of 75 mN. **(g)** Spike frequencies of SAMs in response to a series of increasing displacement stimuli. Mean values ± SEM. **(h)** Mechanical thresholds, the minimum force needed to evoke an action potential, of SAMs. Individual data points and mean values ± SEM are shown. Data was analysed using a one-way ANOVA. **(i)** Example traces of D-hair receptor responses to a ramp-and-hold stimulus with a probe velocity of 0.45 mm/s. **(j)** Example traces of AM responses to a ramp-and-hold stimulus with an indentation force of 125 mN. Abbreviations: SAM, slowly-adapting mechanoreceptor; RAM, rapidly-adapting mechanoreceptor; D-hair, Down-hair; AM, A-mechanonociceptor. Statistical differences calculated with a two-way repeated measures ANOVA (Bonferroni post hoc test).

### STOML3 is required for muscle afferents to acquire mechanosensitivity in the skin

In *stoml3* mutant mice, mechanosensitive Aβ-fibres, including SAMs and RAMs (Figure 3a), and AM fibres (Figure 3b) were found in all three experimental groups (‘intact’, ‘self’, and ‘cross’). As in wild-type mice D-hair receptors were only recorded in the intact and self-anastomosed sural nerves (Figure 3b). In addition, we found tap-units in all three preparations. These were afferents which only fire one spike to extremely rapid high amplitude mechanical stimulation (Figure 3a). Such units were commonly encountered in *stoml3* mutant mice in our previous studies (Wetzel et al., 2007). It was immediately obvious that it was very difficult to find mechanosensitive afferent fibres in the cross-anastomosed gastrocnemius nerve, indeed most fibres with a receptive field were found to be so-called tap-units (Figure 3a-e). Considering the sparsity of responsive fibres found in the cross-anastomosed nerve in *stoml3* mutant mice, we employed an electrical search protocol to assess the proportion of fibres with apparently no mechanosensitive receptive field in the cross-anastomosed gastrocnemius nerve. We found a dramatic and statistically significant increase in the proportion of mechano-insensitive Aβ-fibres and Aδ-fibres in the cross-anastomosed gastrocnemius nerve innervating the skin in *stoml3* mutants compared to controls (Aδ-fibres – C57BL/6: 26% (4/22 fibres), n = 4 mice; *Stoml3^-/-^*: 62% (25/41 fibres), n = 3 mice; two-sided Fisher’s exact test: p = 0.001; Aβ-fibres – C57BL/6: 18% (10/38 fibres), n = 4 mice; *stoml3^-/-^*: 61% (33/53 fibres), n = 3 mice; two-sided Fisher’s exact test: p = 0.001; Figure 3c). Of the remaining mechanosensitive Aβ-fibres in *stoml3* mutant mice (~39% of the total fibres) more than half (55%) were classified as tap-units in the cross-anastomosed nerve. This was in marked contrast to wild-type cross-anastomosed nerves in which only 5% of the already large number of mechanosensitive fibres (82% of all fibres) were classified as tap-units. The large increase in the number of tap-units was highly statistically different between wild-type and *stoml3* mutants (C57BL/6: 5% (1/19 fibres), n = 9 mice; *stoml3^-/-^*: 55% (15/27 fibres), n = 10 mice; two-sided Fisher’s exact test: p < 0.0001; Figure 3d,e, example traces are shown). Tap-units clearly represent sensory fibres which would be extremely difficult to activate by natural touch stimuli *in vivo*.

**Figure 3.**
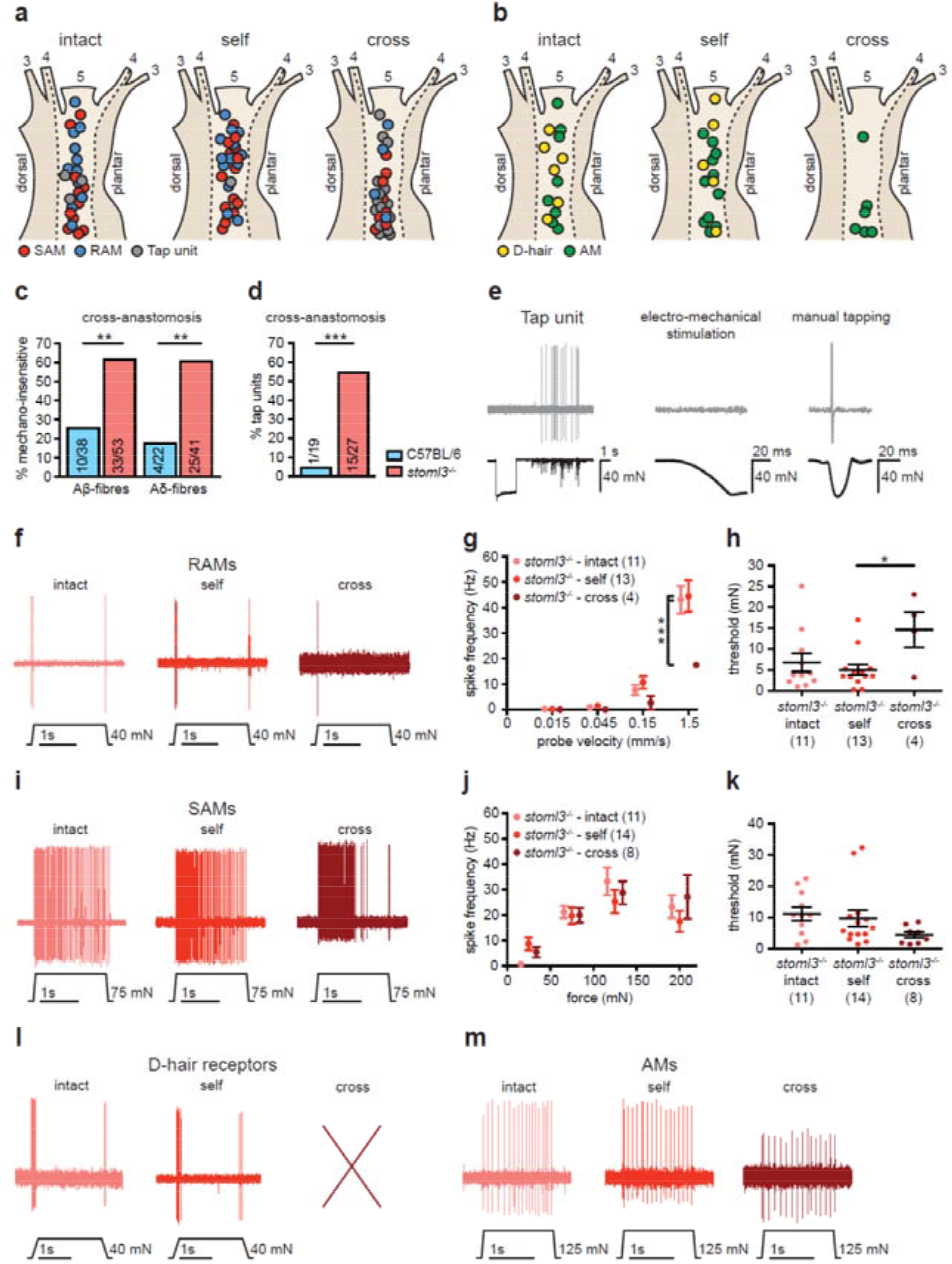
Response properties of muscle afferents newly innervating the skin compared to intact and regenerated cutaneous afferents in *stoml3* mutant mice. **(a,b)** Receptive field locations of intact cutaneous, regenerated cutaneous, and redirected muscle **(a)** Aβ-fibres (SAMs, red; RAMs, blue; tap-units, grey) and **(b)** Aδ-fibres (D-hair receptors, yellow; AMs, green). **(c)** Proportions of mechano-insensitive Aβ- and Aδ-fibres in the skin territory of the sural nerve innervated by redirected muscle afferents in *stoml3* mutant (red, n = 3) as compared to control (blue, n = 4) mice. **(d)** Proportion of units exhibiting tap-unit responses in skin innervated by redirected muscle afferents in *stoml3* mutant (red, n = 10) compared to control (blue, n = 9) mice. **(e)** Example trace of a tap-unit only responding to manually delivered brisk tapping, but not to a controlled stimulus delivered by mechanoelectrical stimulator. **(f)** Example traces of RAM responses to a ramp-and-hold stimulus with a probe velocity of 1.5 mm/s. **(g)** Spike frequencies of RAMs in response to ramp-and-hold stimuli with increasing ramp velocities. Mean values ± SEM Statistical differences calculated with a two-way repeated measures ANOVA (Bonferroni post hoc test). **(h)** Mechanical thresholds of RAMs measured in response to a sinusoidal vibration stimulus (50 Hz). Individual data points and mean values ± SEM. Statistical analysis with a one-way ANOVA (Tukey’s multiple comparison test). **(i)** Example traces of SAM responses to a ramp-and-hold stimulus, 75 mN indentation force **(j)** Spike frequencies of SAMs in response to a series of increasing displacement stimuli. Mean values ± SEM. **(k)** Mechanical thresholds for SAMs. Individual data points and mean values ± SEM. Statistical calculated with a one-way ANOVA. **(l)** Example traces of D-hair receptor responses to a ramp-and-hold stimulus with a probe velocity of 0.45 mm/s. **(m)** Example traces of AM responses to a ramp-and-hold stimulus, 125 mN indentation force. Abbreviations: SAM, slowly-adapting mechanoreceptor; RAM, rapidly-adapting mechanoreceptor; D-hair, Down-hair; AM, A-mechanonociceptor.

We next assessed the stimulus-response properties of the remaining mechanosensitive Aβ-mechanoreceptors in the cross-anastomosed gastrocnemius nerve (Table 2; Figure 3f,i, example traces are shown), which we estimated to be around 18% of all fibres in *stoml3* mutants compared to 78% in controls. The suprathreshold responses to moving stimuli of RAMs found in the cross-anastomosed gastrocnemius nerve preparation were significantly decreased compared to RAMs found in the intact or self-anastomosed nerve, and this was particularly prominent for the fastest stimulus of 1.5 mm/s (‘intact’: n = 11 fibres; ‘self’: n = 13 fibres; ‘cross’: n = 4 fibres; two-way repeated measures ANOVA: F(6,75) = 3.136, p = 0.009; Bonferroni post hoc test; Figure 3g). In addition, mechanical thresholds of RAMs were found to be significantly higher in cross-anastomosed muscle nerves compared to self-anastomosed nerves (‘intact’: 6.8 ± 2.2 mN, n = 11 fibres; ‘self’: 5.0 ± 1.3 mN, n = 13 fibres; ‘cross’: 14.6 ± 4.2 mN, n = 4 fibres; one-way ANOVA: F(2,25) = 3.587, p = 0.043; Tukey’s multiple comparison test; Figure 3h). Interestingly the few sensory afferents found with properties of SAMs in the cross-anastomosed gastrocnemius nerve had stimulus-response properties and mechanical thresholds that were indistinguishable from those in intact or self-anastomosed nerves (stimulus-response function – ‘intact’: n = 11 fibres; ‘self’: n = 14 fibres; ‘cross’: n = 8 fibres; two-way repeated measures ANOVA: F(6,90) = 1.532, p = 0.177; Figure 3j; mechanical thresholds – ‘intact’: 11.2 ± 2.1 mN, n = 11 fibres; ‘self’: 9.7 ± 2.6 mN, n = 14 fibres; ‘cross’: 4.5 ± 1.0 mN, n = 8 fibres; one-way ANOVA: F(2,30) = 1.894, p = 0.009; Figure 3k).

Unlike in wild-type mice, D-hair receptors found in the self-anastomosed *stoml3* mutant sural nerve fired significantly fewer spikes to ramp-and-hold stimuli compared to D-hair receptors in the intact *stoml3* mutant sural nerve (Figure 3l, examples traces are shown) (‘intact’: n = 6 fibres; ‘self’: n = 5 fibres; two-way repeated measures ANOVA: F(3,27) = 6.821, p = 0.001; Bonferroni post hoc test; Figure 3, Figure 3-1). However, there was no difference in mechanical thresholds between D-hair receptors found in *stoml3* mutant mice within the intact or self-anastomosed nerve (‘intact’: 0.7 ± 0.4 mN, n = 6 fibres; ‘self’: 0.6 ± 0.2 mN, n = 5 fibres; two-tailed unpaired t-test: t(9) = 0.1511, p = 0.883; Figure 3, Figure 3-1b). Thinly myelinated nociceptors or AMs recorded from *stoml3* mutants (Figure 3m, example traces are shown) exhibited similar stimulus-response functions and mechanical thresholds in intact, self-, and cross-anastomosed nerves (stimulus-response function – ‘intact’: n = 8 fibres; ‘self’: n = 11 fibres; ‘cross’: n = 6 fibres; two-way repeated measures ANOVA: F(6,66) = 1.000, p = 0.433; Figure 3, Figure 3-1c; mechanical thresholds – ‘intact’: 54.8 ± 7.5 mN, n = 8 fibres; ‘self’: 39.9 ± 8.5 mN, n = 11 fibres; ‘cross’: 32.2 ± 5.3 mN, n = 6 fibres; one-way ANOVA: F(2,22) = 1.755, p = 0.196; Figure 3, Figure 3-1d).

The striking lack of mechanosensitive fibres in the cross-anastomosed gastrocnemius nerve may have been due to an inability of sensory fibres to regenerate and form appropriate endings in *stoml3* mutant mice. We used transmission electron microscopy to quantify the number of myelinated axons that regenerated distal to the cross-anastomosis site in wild-type and *stoml3* mutant mice (Figure 4a, n = 3 mice each, representative images are shown). We found equal numbers of regenerated axons in both genotypes (C57BL/6 ‘cross’: 722.4 ± 133.9, n = 3 mice; *stoml3^-/-^* ‘cross’: 649.2 ± 54.92, n = 3 mice; two-tailed unpaired t-test: t(4) = 0.5061, p = 0.6394; Figure 4b). We further assessed the innervation of the skin by redirected muscle afferents in both controls and *stoml3* mutants. Using immunocytochemistry to label all sensory fibres with antibodies against protein gene product 9.5 (PGP9.5) or myelinated sensory fibres with antibodies against neurofilament 200 (NF200) we could show that hair follicles were innervated by muscle sensory afferents (n = 3 mice each; Figure 4c, representative images are shown). Thus, lanceolate endings positive and negative for neurofilament 200 were found in the skin innervated by the muscle nerve in both wild-type and *stoml3* mutant mice. Furthermore, these endings were similar to those found in the intact sural nerve territory. We also labelled the endings of putative SAMs in the skin using antibodies against cytokeratin 8/18 (TROMA-I) to label Merkel cells and found that these cells were innervated by muscle sensory axons positive for NF200 in both wild-type and *stoml3* mutant mice (n = 3 mice each; Figure 4c, representative images are shown). We conclude that the remarkable ability of muscle afferents to form sensory endings appropriate for the skin does not in fact depend on the presence of STOML3.

**Figure 4.**
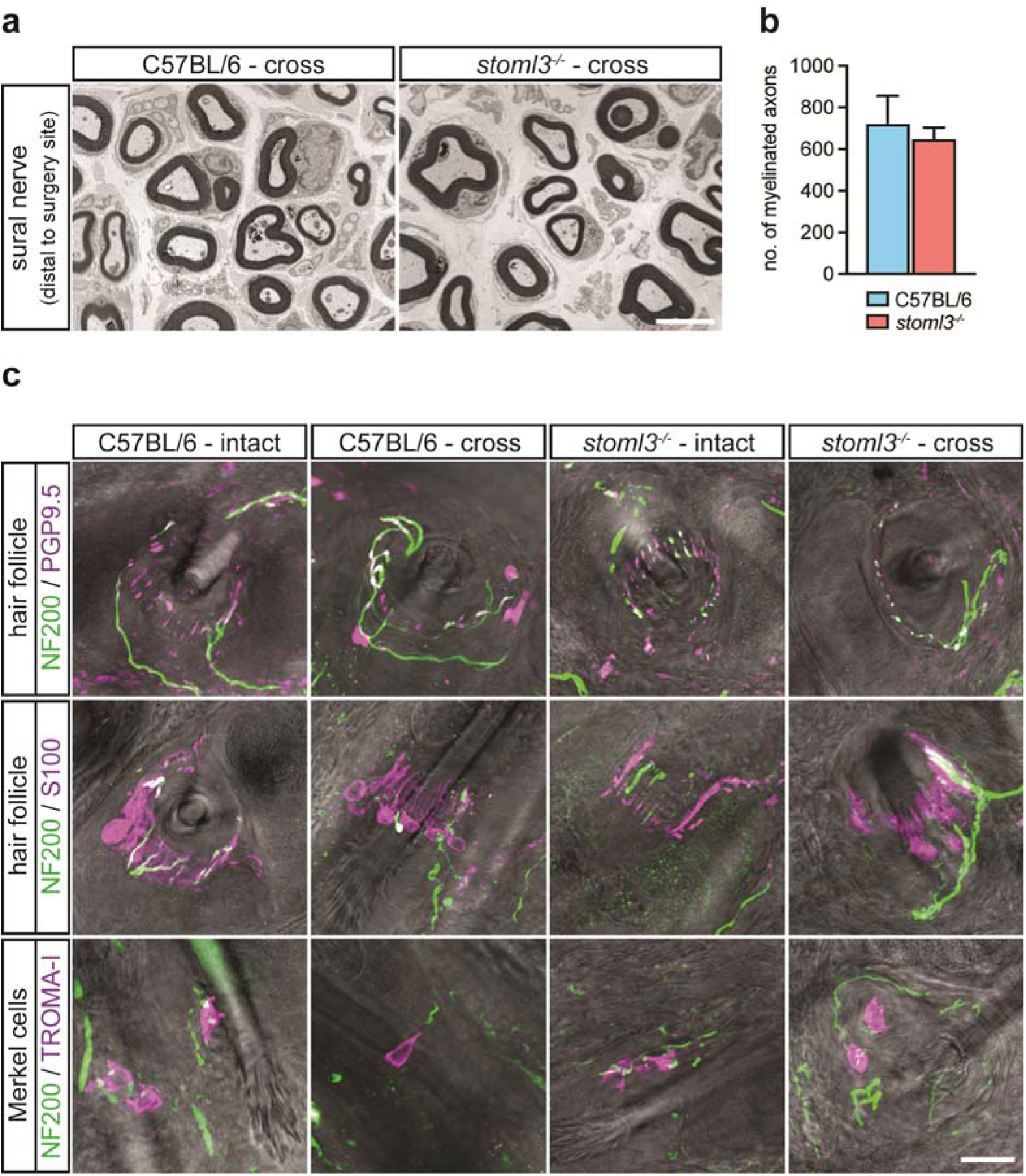
Skin innervation by muscle afferents in *stoml3* mutant and control mice. **(a)** Transverse electron microscopic images of cross-anastomosed gastrocnemius nerves innervating the skin distal to the surgery site in *stoml3* mutant and wild-type mice. Scale bar: 3 μm. **(b)** Numbers of myelinated fibres counted in cross-anastomosed sural nerves distal to the surgery site in *stoml3* mutant (red, n = 3) as compared to control (n = 3, blue) mice. Mean values ± SEM are shown. Data was analysed using a two-tailed unpaired t-test. **(c)** Fluorescent images of hair follicles and Merkel cells in the skin territory of the sural nerve innervated by intact sural nerve afferents and redirected muscle afferents. Scale bar: 20 μm. Abbreviations: NF200, neurofilament 200; PGP9.5, protein gene product 9.5; TROMA-I, trophectodermal monoclonal antibody against cytokeratin 8.

### Somatotopic map formation is blurred in *stoml3* mutant mice

Previous studies in the rat have shown a remarkable amount of functional plasticity of muscle afferents redirected to the skin. Redirected muscle afferents engage new reflexes and make new synaptic connections with dorsal horn neurons in a somatotopically appropriate manner (McMahon and Wall, 1989; Lewin and McMahon, 1993). In order to study structural plasticity of sensory afferents after regeneration we established a quantitative method to study somatotopic mapping of sensory afferent terminals in the spinal cord (Tröster et al., 2018). We first used this tracing methodology to map the accuracy of sensory afferent projections in touch-deficient *stoml3* mutant mice as the presence of deficits in the intact condition could have a bearing on what happens after nerve regeneration. Sensory afferents innervating the second and third digit of the left and right hind paw, respectively, were labelled using subcutaneous injections of cholera toxin subunit B conjugated with Alexa Fluor^®^ 594 (CTB) which is selectively endocytosed by myelinated fibres (Wan et al., 1982; Robertson and Arvidsson, 1985). Five days after the injection the central terminal fields of cutaneous fibres innervating the skin of the second and third digit of the left and right hind paw, respectively, were mapped in their entirety in un-sectioned cleared spinal cords of four-week old *stoml3* mutant and control mice. Control mice were the C57BL/6 strain as the *stoml3* mutant line had been back-crossed onto the same background for at least 10 generations. To visualize CTB-labelled projections we evaluated several of the published optical clearing methods (Staudt et al., 2007; Kloepper et al., 2010; Ertürk et al., 2011, 2012; Costantini et al., 2015) and found that immersion clearing using 2’2-thiodiethanol (TDE) produced the least tissue shrinkage whilst providing sufficient imaging depth and preserving fluorescence intensity (Figure 5-1).

**Figure 5.**
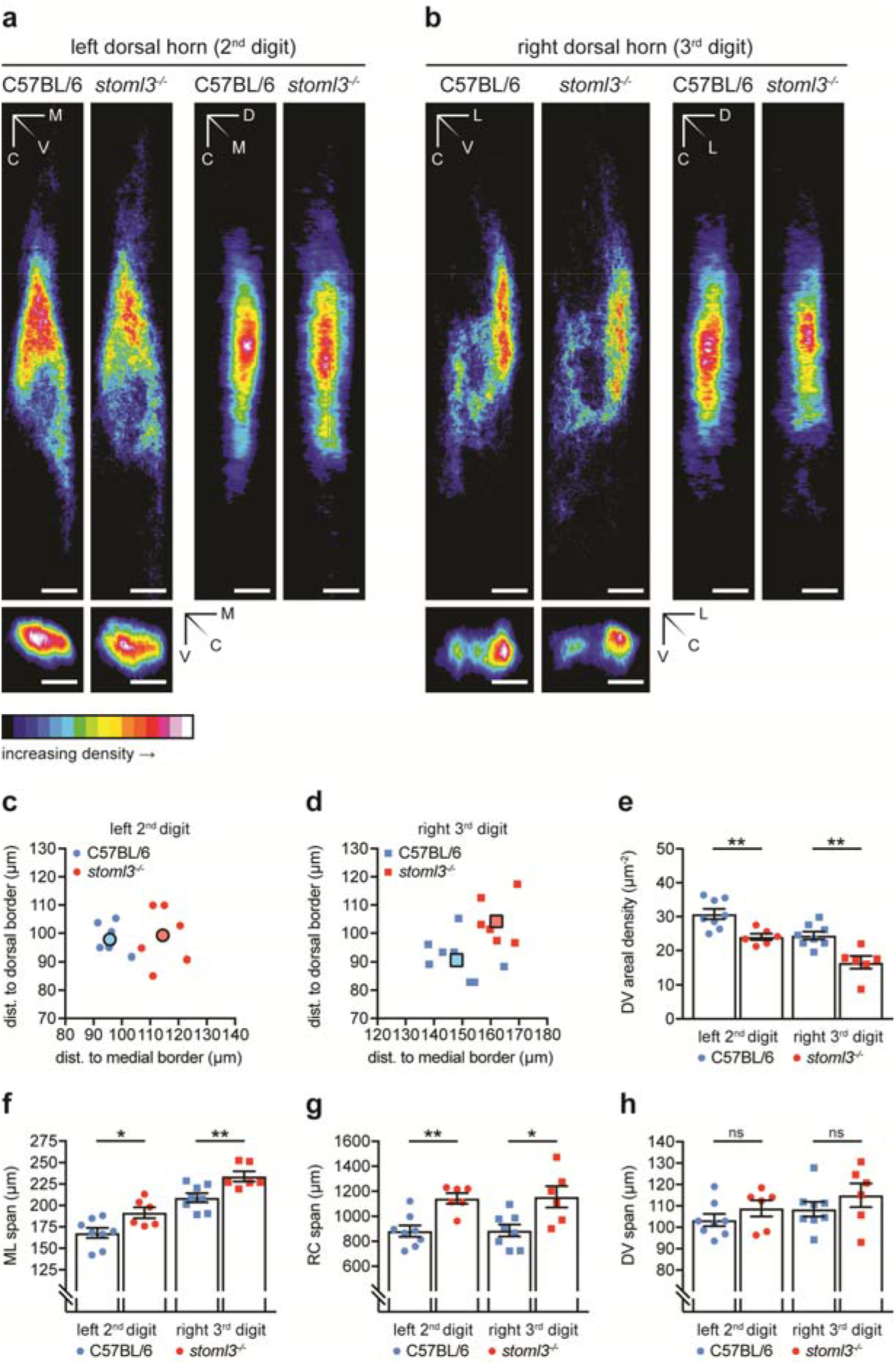
Morphometric and density measurements of spinal terminal fields of fibres innervating the left second and right third hind paw digit in *stoml3* mutant and control mice. **(a,b)** Averaged summed dorso-ventral, medio-lateral, and rostro-caudal projections of spinal terminal fields of fibres innervating **(a)** the left second and **(b)** right third hind paw digit in *stoml3* mutant (n = 6) and control (n = 8) mice. The ImageJ colour lookup table “16 Colors” was applied. The colour code in each pixel denotes the number of voxels found in corresponding positions along different axes averaged across mice. Scale bars: 100 μm. **(c,d)** The locations of terminal field foci of fibres innervating **(c)** the second digit of the left hind paw (circles) and **(d)** the third digit of the right hind paw (squares) relative to the medial and dorsal grey/white matter border in *stoml3* mutant (red, n = 6) and control (blue; n = 8) mice. Individual data points are shown. **(f,g,h)** Spans of terminal fields of fibres innervating the left second (circles) and right third (squares) hind paw digit in the **(f)** medio-lateral (ML), **(g)** rostro-caudal (RC), and **(h)** dorso-ventral (DV) dimension in *stoml3* mutant (red, n = 6) and control (blue, n = 8) mice. Individual data points and mean values ± SEM are shown. Each data set was compared using a two-tailed unpaired t-test. **(e)** Areal densities of terminal fields innervating the left second (circles) and right third (squares) hind paw digit in dorso-ventral projections in *stoml3* mutant (red, n = 6) and control (blue, n = 8) mice. Individual data points and mean values ± SEM are shown. Each data set was compared using a two-tailed unpaired t-test. Abbreviations: M, medial; L, lateral; D, dorsal; V, ventral; C, caudal; ML, medio-lateral; RC, rostro-caudal; DV, dorso-ventral.

We performed experiments to ensure that a comparative analysis of CTB-labelled spinal terminal fields could be made between *stoml3* mutants and controls. Importantly, the medio-lateral widths of spinal cord dorsal horns, measured at a depth of 80 μm from the dorsal surface, were not different in *stoml3* mutants compared to controls (C57BL/6: 1091 ± 20 μm, n = 8 mice; *stoml3^-/-^*: 1114 ± 16 μm, n = 6 mice; two-tailed unpaired t-test: t(12) = 0.8819, p = 0.395; Figure 5, Figure 5-2a). Furthermore, reliable and consistent numbers of sensory neurons were labelled as demonstrated by counting total numbers of CTB-labelled neurons in lumbar DRGs 3, 4, and 5 that innervate the hind limb skin; on average there was no difference in the numbers of labelled sensory neurons between genotypes. (second left hind paw digit – C57BL6: 124.2 ± 7.8, n = 5 mice; *stoml3^-/-^*: 127.6 ± 8.6, n = 5 mice; two-tailed unpaired t-test: t(8) = 0.2915, p = 0.778; third right hind paw digit – C57BL6: 132.6 ± 13.2, n = 5 mice; *stoml3^-/-^*: 129.6 ± 3.9, n = 5 mice; two-tailed unpaired t-test with Welch’s correction: t(4.691) = 0.2187, p = 0.836; Figure 5-2b). Furthermore, CTB-labelling was restricted to the same skin areas in both control and *stoml3* mutant mice (n = 3 mice each; Figure 5-2c, representative images shown).

The segmental and laminar location of CTB-labelled terminals, as well as their overall geometry were similar between genotypes (Figure 5a,b). For quantitative analyses the three-dimensional centres of mass of the voxel clouds representing CTB-labelled fibre terminals in binary image stacks were measured and the terminal fields were aligned to their centre of mass (Figure 5, Figure 5-3 Video 5-1). Summed dorso-ventral, rostro-caudal, and/or medio-lateral projections of the binary image stacks were constructed and a colour lookup table was applied to enable visualisation of terminal fields in *stoml3* mutant and control mice (Figure 5a,b; Supplementary Video 1). First, we determined the locations of the terminal field foci relative to the medial and dorsal grey/white matter border in *stoml3* mutant and control mice. The foci of terminal fields of fibres innervating the second digit were on average shifted laterally by 18.75 μm and ventrally by 1.42 μm in *stoml3* mutant mice compared to controls (Figure 5c) and the foci of labelled terminal fields representing the third digit were on average shifted laterally by 14.00 μm and ventrally by 13.54 μm in *stoml3* mutant mice compared to controls (Figure 5d). In the two-dimensional space the foci of the terminals fields of fibres innervating the second and third digit were on average shifted by 18.8 μm and 19.5 μm, respectively, in *stoml3* mutant mice compared to controls (Figure 5c,d). The terminal field foci tended to be shifted in a rostro-lateral direction as can be seen by examining the summed termination fields (Figure 5a,b). Next, we determined the maximal extent of the terminal fields in the medio-lateral (ML), rostro-caudal (RC), and dorso-ventral (DV) dimension by measuring the dimensions of the minimum bounding rectangle that enclosed all pixels in dorso-ventral and rostro-caudal summed projections of the binary image stacks. The extent of the terminal fields in *stoml3* mutant and control mice were found to be significantly different (Table 2; Figure 5f-h). The spinal terminal fields of fibres innervating the second and third digit of the left and right hind paw, respectively, extended 14% and 12% further in the medio-lateral dimension (Table 2; Figure 2f) and 30% in the rostro-caudal dimension in *stoml3* mutant mice compared to control mice (Table 1; Figure 2g). No differences in the extent of terminal fields in the dorso-ventral dimension were observed between genotypes (Table 2; Figure 2h). Despite the fact that terminal fields were expanded in *stoml3* mutants the numbers of voxels representing CTB-labelling was not different between the two genotypes (left terminal field – C57BL/6: 3.14 ± 0.418 × 10^6^, n = 8 mice; *stoml3^-/-^*: 2.78 ± 0.183 × 10^6^, n = 6 mice; two-tailed unpaired t-test with Welch’s correction: t(9.457) = 0.792, p = 0.448; right terminal field – C57BL/6: 2.30 ± 0.271 × 10^6^, n = 8 mice; *stoml3^-/-^*: 1.79 ± 0.260 × 10^6^, n = 6 mice; two-tailed unpaired t-test: t(12) = 1.319, p = 0.212; Figure 5, Figure 5-3). The lack of change in voxel numbers led us to suspect that there was a decrease in the density of the spinal terminal fields in *stoml3* mutants compared to controls which was already apparent in the summed intensity projections shown in Figure 5a,b. We calculated the areal density of the terminal fields separately in the dorso-ventral, medio-lateral, and rostro-caudal summed projections by dividing the total number of voxels in binary image stacks by the area occupied by pixels in each of the three projections (Figure 5, Figure 5-3). The areal densities of terminal fields of fibres innervating the second and third digits, respectively, were significantly lower in all three projections in *stoml3* mutants compared to control mice (Figure 5e; Figure 5-3b-d). In dorso-ventral projections, the areal density of fibres innervating the hind paw digits were between 22-32% less dense in *stoml3* mutants compared to controls and this was statistically significant (left terminal field – C57BL/6: 30.8 ± 1.5 voxels/μm^2^, n = 8 mice; *stoml3^-/-^*: 24.0 ± 0.9 voxels/μm^2^, n = 6 mice; two-tailed unpaired t-test: t(12) = 3.463, p = 0.005; right terminal field – C57BL/6: 24.4 ± 1.2 voxels/μm^2^, n = 8 mice; *stoml3^-/-^*: 16.5 ± 1.8 voxels/μm^2^, n = 6 mice; two-tailed unpaired t-test: t(12) = 3.848, p = 0.002; Figure 5e). Our quantitative analysis demonstrates that afferent terminal fields in the deep dorsal horn occupy a larger area of the spinal cord and are less dense in *stoml3* mutant mice compared to controls.

### Mechanosensory silence does not prevent structural plasticity

We next asked whether we could use CTB-labelling to visualise the structural plasticity of muscle afferents in the dorsal horn. Normally myelinated sensory fibres from the gastrocnemius muscle (proprioceptors and myelinated muscle nociceptors) terminate in deeper dorsal laminae and predominantly in the ventral horn (Brown and Fyffe, 1978, 1979; Brown, 1981), whereas skin mechanoreceptors project to (inter-)neurons in laminae III to V in a somatotopically organized fashion (Brown and Culberson, 1981; Shortland et al., 1989; Shortland and Woolf, 1993). We labelled axons within the regenerated nerve by injecting 2 μl CTB into the nerve distal to the anastomosis site and waited 5 days before visualising the transganglionically transported tracer in the dorsal horn (Belyantseva and Lewin, 1999). We could reliably trace the central terminals of intact and regenerated sural nerve fibres as well as cross-anastomosed muscle afferents newly innervating the skin in both wild-type and *stoml3* mutant mice (Figure 6). The tributary branches of the sciatic nerve, namely the cross-anastomosed gastrocnemius and sural nerves, the common peroneal nerve, and the tibial nerve were microscopically examined for CTB-labelling to ensure restricted labelling of fibres running in the cross-anastomosed gastrocnemius nerve. Only the sural nerve trunk containing redirected gastrocnemius sensory fibres was labelled (n = 3 mice each; Figure 6, Figure 6-1, representative images are shown).

**Figure 6.**
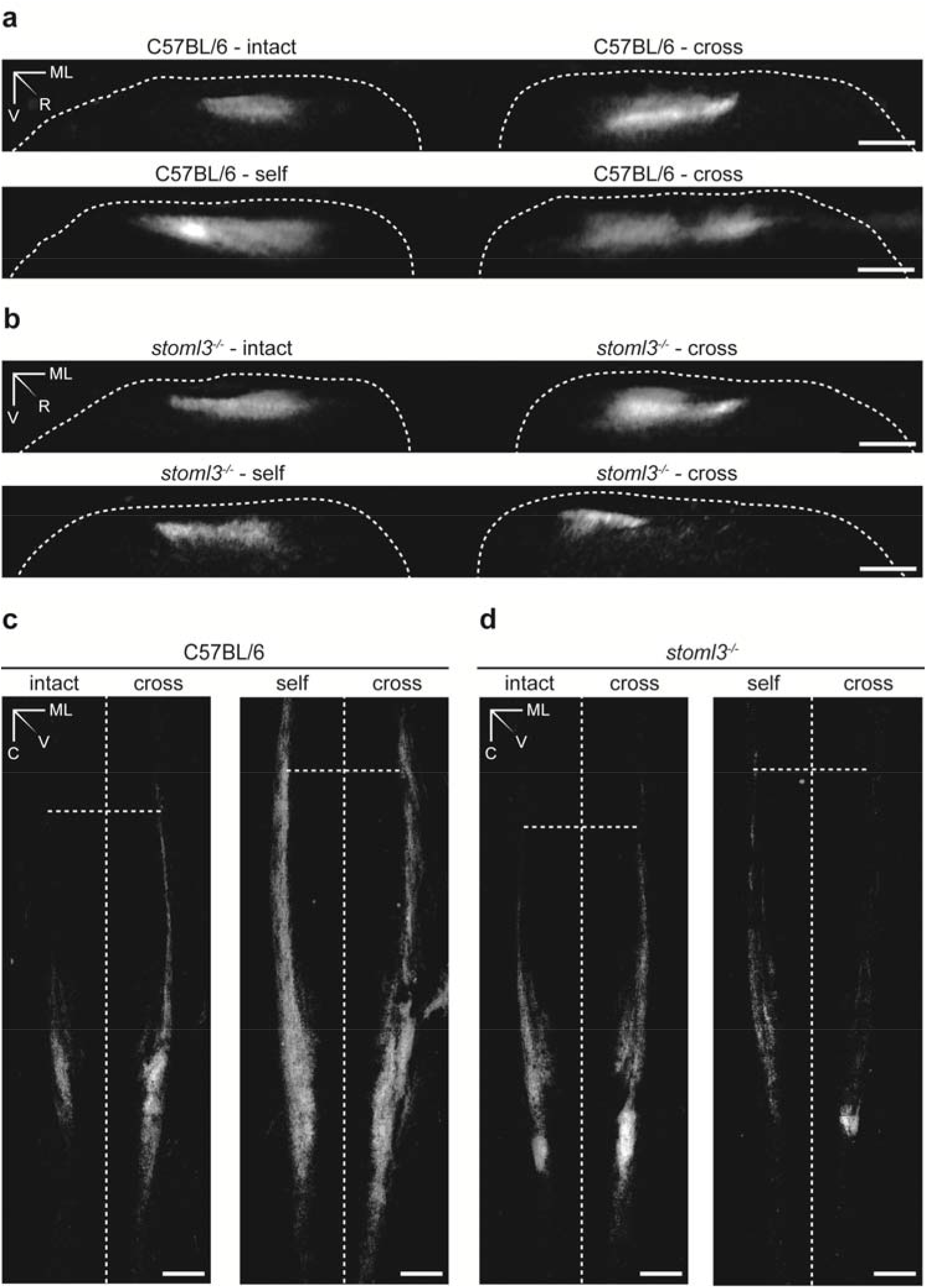
Spinal terminal fields of intact, self-anastomosed, and cross-anastomosed nerve afferents innervating the sural nerve skin territory in wild-type and *stoml3* mutant mice. **(a,b)** Summed rostro-caudal projections of terminal fields of muscle afferents redirected towards the skin in comparison to intact and regenerated cutaneous afferents in **(a)** wild-type and **(b)** *stoml3* mutant mice. The dorsal grey/white matter border is marked with a white dashed line. Scale bars: 100 μm. **(c,d)** Summed dorso-ventral projections of terminal fields innervating of muscle afferents redirected towards the skin in comparison to intact and regenerated cutaneous afferents in **(c)** wild-type mice and **(d)** *stoml3* mutant mice. Vertical dashed lines mark the posterior median sulcus, horizontal dashed lines mark the border between spinal lumbar segments 3 and 4. Scale bars: 200 μm. Abbreviations: ML, medio-lateral; R, rostral; V, ventral; C, caudal.

The raw images of CTB-labelled terminals in the spinal cord were binarised, and summed dorso-ventral and rostro-caudal projections were generated to assess the laminar positioning and the somatotopic arrangement of the terminal fields. As shown in Figure 6a (representative images are shown), the central terminals of muscle afferents newly innervating the skin terminated at the same dorso-ventral level as cutaneous afferents of the intact and self-anastomosed sural nerve (‘intact’ and ‘self’: n = 3 mice each, ‘cross’: n = 7 mice; Figure 6a, representative images are shown). Furthermore, the central terminals of muscle afferents established somatotopically organized projections resembling the terminal fields of intact and self-anastomosed sural nerves (‘intact’ and ‘self’: n = 3 mice each, ‘cross’: n = 7 mice; Figure 6c, representative images are shown). Injections of CTB into the popliteal fossa of mice in which the sural nerve was transected did not lead to labelling of central afferents. Surprisingly, spinal terminals of muscle afferents now innervating the skin in *stoml3* mutant mice reorganized to terminate in a somatotopically organized fashion in dorsal horn laminae comparable to those of the intact and regenerated sural nerve (‘intact’ and ‘self’: n = 3 mice each, ‘cross’: n = 6 mice; Figure 6b,d, representative images are shown). Thus, we have demonstrated that muscle afferents confronted with a new target in the skin can exhibit substantial structural plasticity in that they form new anatomical connections with somatotopically appropriate dorsal horn neurons. Strikingly, a substantial loss of mechanosensitivity in most of the redirected gastrocnemius afferents in the skin of *stoml3* mutant mice does not prevent these afferents from displaying similar structural plasticity to controls.

## Discussion

It has known for some time that regenerating axons efficiently regain receptor properties in their new target after nerve transection (Burgess and Horch, 1973; Fawcett and Keynes, 1990). However, to date nothing was known about the molecular factors that are required for the re-acquisition of a mechanosensitive receptive fields after regeneration. In contrast to wild-type mice, where virtually all muscle afferents can make mechanosensitive endings in the skin (Figure 2), only a small fraction (<20%) of muscle afferents from *stoml3* mutants were capable of acquiring normal mechanosensitivity when directed to the skin (Figure 3). Only muscle afferents that acquired the properties of slowly adapting mechanoreceptors (SAMs) showed mechanosensitivity similar to controls (Figure 3). We presume that these muscle afferents form peripheral endings associated with Merkel cells (Figure 4), which themselves contribute to SAM mechanosensitivity (Maksimovic et al., 2014). Thus, *stoml3* appears to be genetically required for most muscle afferents to form mechanosensitive endings in the skin. Despite the lack of mechanosensory function, muscle afferents formed morphological end-organs in the skin appropriate to the new target in the absence of *stoml3* (Figure 4). Thus, the STOML3 protein is dispensable for the formation of end-organ morphology, but is still required for most muscle afferents to acquire mechanosensitivity in the skin.

Following re-routing to skin muscle afferents display remarkable functional plasticity in the spinal cord (McMahon and Wall, 1989; Lewin and McMahon, 1993), forming somatotopically appropriate connections on to dorsal horn neurons which normally receive little synaptic input from intact muscle afferents (Lewin and McMahon, 1993). Here we show that there is a substantial anatomical rearrangement of the central terminals of myelinated muscle afferents after re-routing to skin. The synaptic terminals of muscle afferents innervating the skin could be robustly visualized after CTB-tracing in a restricted region of the dorsal horn that corresponds to the appropriate somatotopic territory occupied by afferents from the intact or self-anastomosed sural nerve (Figure 6). This anatomical plasticity was robust and was observed in all animals studied, including *Stom3^-^* mutant mice. Muscle afferents do not normally project to the same region of the dorsal horn before regeneration, but this was difficult to show directly. We carried out CTB-tracing experiments from the intact gastrocnemius nerve, but never observed any signal in the cleared spinal cord after two photon imaging. It was impossible to tell under these circumstances whether the tracing had failed (a rare occurrence in our hands) or whether as previously documented the muscle afferent synapses in the dorsal horn are so sparse (Molander and Grant, 1987; Hoheisel et al., 1989; Panneton et al., 2005) that labelling was not detectable in the cleared tissue. This striking structural plasticity also occurred in *stoml3* mutant mice despite an almost complete loss of mechanosensitivity of muscle afferents innervating the skin (Figure 3). We could not detect any major difference in the central projection of afferents from muscle nerves innervating skin between wild-type and *stoml3* mutant^-^ mice. However, the variability in the central projections of cross-anastomosed muscle afferents between animals made it impossible to reliably quantify differences in projection patterns between genotypes. If activity arising from the periphery plays a role it could be that the reduced mechanically evoked activity in *stoml3* mutant mice is still sufficient over time to direct anatomical plasticity.

Using a precise and unbiased method to reconstruct the somatotopy of cutaneous projections after CTB-tracing we observed more diffuse and dispersed representation of the skin in the spinal cord of *stoml3* mutant mice (Figure 5). Around 40% of mechanoreceptors in *stoml3* mutant mice are mechanically silent (Wetzel et al., 2007, 2017) and we speculate that the lack of stimulus evoked activity may have impaired activity-dependent sharpening of the somatotopic map in these animals (Beggs et al., 2002; Granmo et al., 2008). Indeed, the diffuse somatotopic map that we observed here may be a major reason for reduced tactile acuity in *stoml3* mutant mice (Wetzel et al., 2007). In these experiments we found that each digit is represented within a rostro-caudal band which at its narrowest has a width of less than 100 μm (Figure 5), considering the substantial expansion (up to 30%) of the terminal fields in *stoml3* mutants it is clear that the representation of the digits will overlap. Unfortunately, we could not directly visualise such an overlap as the clearing methodology was not compatible with using two different fluorescent CTB-conjugates.

It is well established that after nerve transection sensory axons reach topologically inappropriate positions in the skin and may not reinnervate the same end-organ as before the lesion (Burgess and Horch, 1973; Lewin et al., 1994; Johnson et al., 1995). An extreme case of adult plasticity is when muscle afferents are forced to regenerate inappropriately to skin, a situation that undoubtedly happens following mixed nerve injury in humans (Rbia and Shin, 2017). The only skin receptor type not found in the cross-anastomosed gastrocnemius nerve in both genotypes were D-hair receptors (Figure 2.3). D-hair receptors are the most sensitive type of skin mechanoreceptor which have thinly myelinated axons and are characterized by high expression of the T-type calcium channel Ca_V_3.2 (Shin et al., 2003; Wang and Lewin, 2011; Lechner and Lewin, 2013; Bernal Sierra et al., 2017; Walcher et al., 2018). There is no evidence that the normal muscle is innervated by thinly myelinated low threshold mechanoreceptors (Mense, 1996), which suggests that nociceptors from the muscle can only acquire properties of nociceptors in the skin.

Mutant mice lacking the PIEZO2 modulating protein STOML3 have deficits in tactile acuity (Wetzel et al., 2007), but unlike PIEZO2 deficient humans and mice, do not have proprioceptive deficits (Ranade et al., 2014; Woo et al., 2015; Chesler et al., 2016; Murthy et al., 2018). Thus, STOML3 may not normally be expressed in muscle proprioceptors. Here we find that the presence of STOML3 is actually required for the vast majority of muscle afferents to form mechanosensitive receptive fields in the skin (Figure 3). These data are consistent with the hypothesis that *de novo* expression of STOML3 in muscle afferents innervating the skin is a pre-requisite for mechanosensitivity. Indeed, we have shown that nerve injury alone is sufficient to up-regulate STOML3 protein in sensory neurons (Wetzel et al., 2017), but it is also possible that signals in the skin instruct muscle afferents to express *stoml3*. One example of a peptide factor that has high expression in skin, but low expression in muscle, and can drive spinal plasticity is nerve growth factor (NGF) (Korsching and Thoenen, 1983; Shelton and Reichardt, 1984; Lewin et al., 1992). However, NGF does not upregulate STOML3 expression in the DRG (Wetzel et al., 2017), but could play a role in driving central plasticity (Lewin et al., 1992). The effects of *stoml3* loss of function were highly specific, as muscle afferents were capable of regenerating to the skin and forming morphologically appropriate sensory endings in the skin without *stoml3*. Regeneration was robust in all cases as similar numbers of myelinated gastrocnemius fibres were present in the distal sural nerve stump after cross-anastomosis in wild-type and *stoml3* mutant mice (Figure 4).

Recent work in flies has established a link between mechanotransduction and regeneration (Song et al., 2019). Here we examined the role of the mechanotransduction protein STOML3 in peripheral nerve regeneration. Sensory axons were able to regenerate and form specialized end-organ morphologies in the absence of the STOML3 protein (Figure 4). However, we found that the presence of STOML3 was necessary for muscle afferents to acquire normal mechanosensitivity in the skin. We also found that the somatotopic organization of cutaneous afferents in the dorsal horn is correct, but significantly less focused and precise in *stoml3* mutants compared to controls (Figure 5). Nevertheless, the central terminals of muscle afferents in the *stoml3* mutant mice exhibit dramatic structural plasticity forming somatotopically appropriate terminals even when stimulus evoked activity was greatly attenuated compared to controls. We conclude that there are likely chemical factors in the skin that can induce expression STOML3 in muscle afferents and direct sprouting of their central terminals into somatotopically appropriate areas of the spinal dorsal horn. However, sensory evoked activity even at a low level may still contribute to this plasticity.

## Acknowledgments

The authors gratefully acknowledge the excellent technical assistance of the Advanced Light Microscopy Facility (Dr. Anje Sporbert, Dr. Zoltán Cseresnyés, Dr. Anca Margineanu, and Matthias Richter) and the Electron Microscopy Facility (Dr. Bettina Purfürst, Christina Schiel) of the Max Delbrück Center for Molecular Medicine. The authors thank V Begay, S Chakrabarti and A Barker for critically reading the MS.

## Supplementary Figures and Video

**Supplementary Video 1. Processing of tiled image stacks of CTB-labelled spinal terminal fields. (a)** Raw tiled image stack taken through a TDE-cleared spinal cord. Two channels were recorded to collect both CTB fluorescence (shown) and autofluorescence (not shown). **(b)** By subtracting the autofluorescence channel from the CTB channel, autofluorescence was removed. **(c)** Using stack histogram-based thresholding, the images were binarised. **(d)** Noise was eliminated by removing single voxels. **(e)** A summed dorso-ventral projection was constructed to aid visualisation and analysis. **(f)** The ImageJ colour lookup table ’16 Colors’ was applied to the summed dorso-ventral projection. Scale bar: 150 μm. Abbreviation: LUT, lookup table.

## Extended Data Figures

**Figure 2-1.**
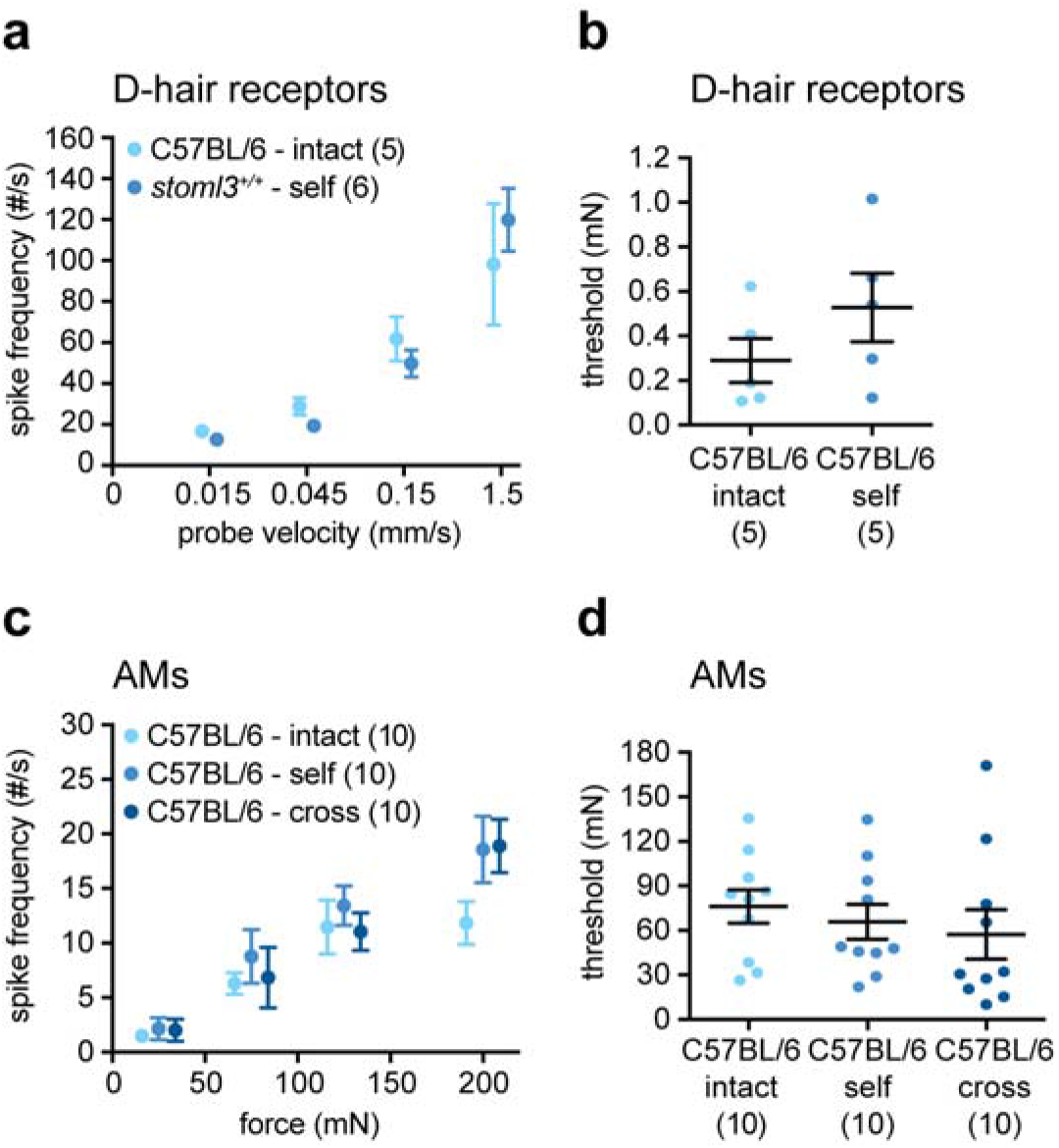
Response properties of muscle Aδ-fibres newly innervating the skin compared to intact and regenerated cutaneous afferents in C57BL/6 mice. **(a)** Spike frequencies in response to ramp-and-hold stimuli with increasing ramp velocities and **(b)** mechanical thresholds measured using a sinusoidal vibration stimulus (25 Hz) of D-hair receptors in C57BL/6 mice. **(c)** Spike frequencies in response to a series of increasing displacement stimuli and **(d)** mechanical thresholds, the minimum force needed to evoke an action potential, of AMs in C57BL/6 mice. Mean values ± SEM or individual data points and mean values ± SEM are shown. Data sets were analysed using two-way repeated measures ANOVAs (Bonferroni post hoc test), one-way ANOVAs, or two-tailed unpaired t-tests. Abbreviations: D-hair, Down-hair; AM, A-mechanonociceptor.

**Figure 3-1.**
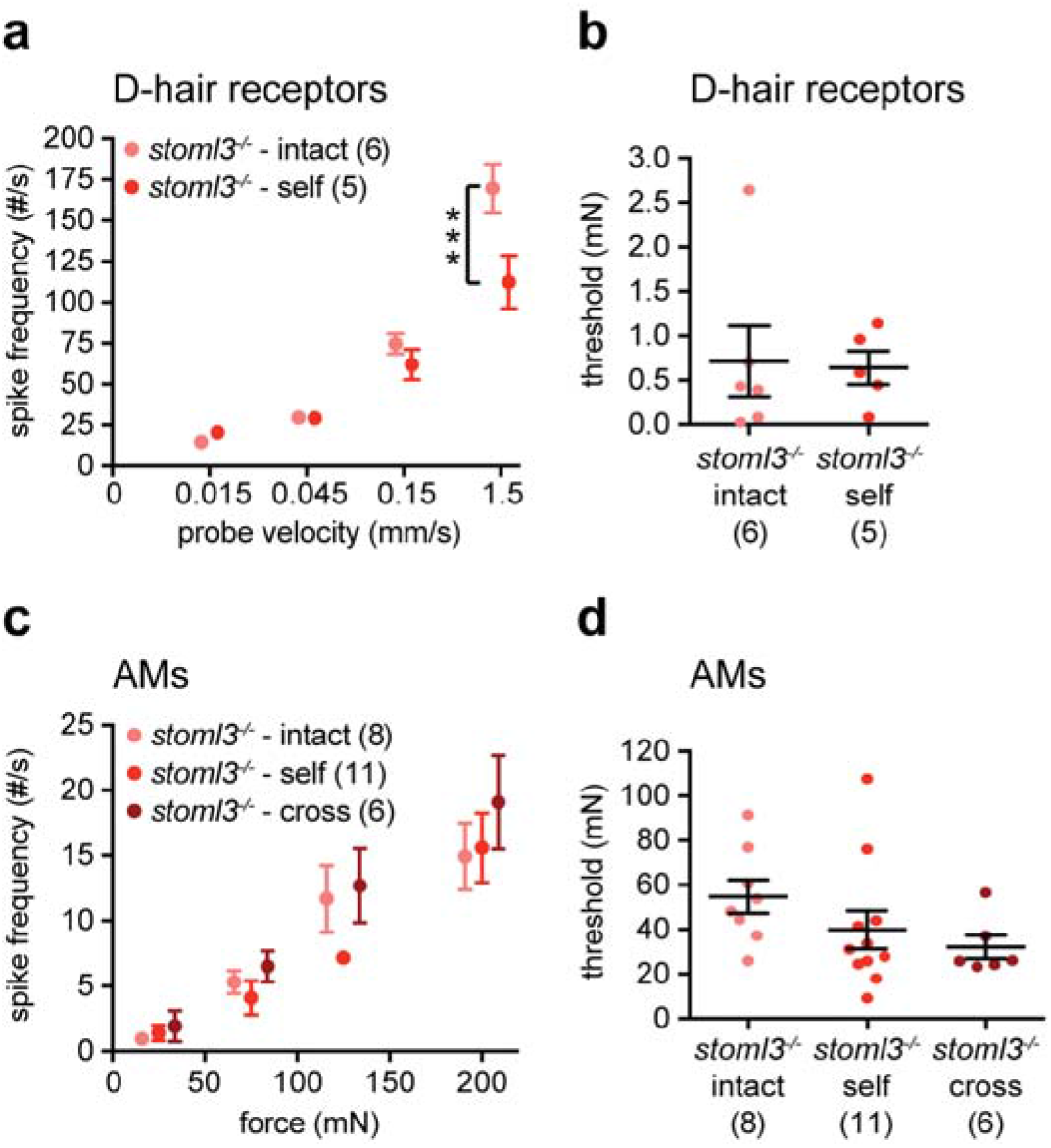
Response properties of muscle Aδ-fibres newly innervating the skin compared to intact and regenerated cutaneous afferents in *stoml3* mutant mice. **(a)** Spike frequencies in response to ramp-and-hold stimuli with increasing ramp velocities and **(b)** mechanical thresholds measured using a sinusoidal vibration stimulus (25 Hz) of D-hair receptors in *stoml3* mutant mice. **(c)** Spike frequencies in response to a series of increasing displacement stimuli and **(d)** mechanical thresholds, the minimum force needed to evoke an action potential, of AMs in *stoml3* mutant mice. Mean values ± SEM or individual data points and mean values ± SEM are shown. Data sets were analysed using two-way repeated measures ANOVAs (Bonferroni post hoc test), one-way ANOVAs, or two-tailed unpaired t-tests. Abbreviations: D-hair, Down-hair; AM, A-mechanonociceptor.

**Figure 5-1.**
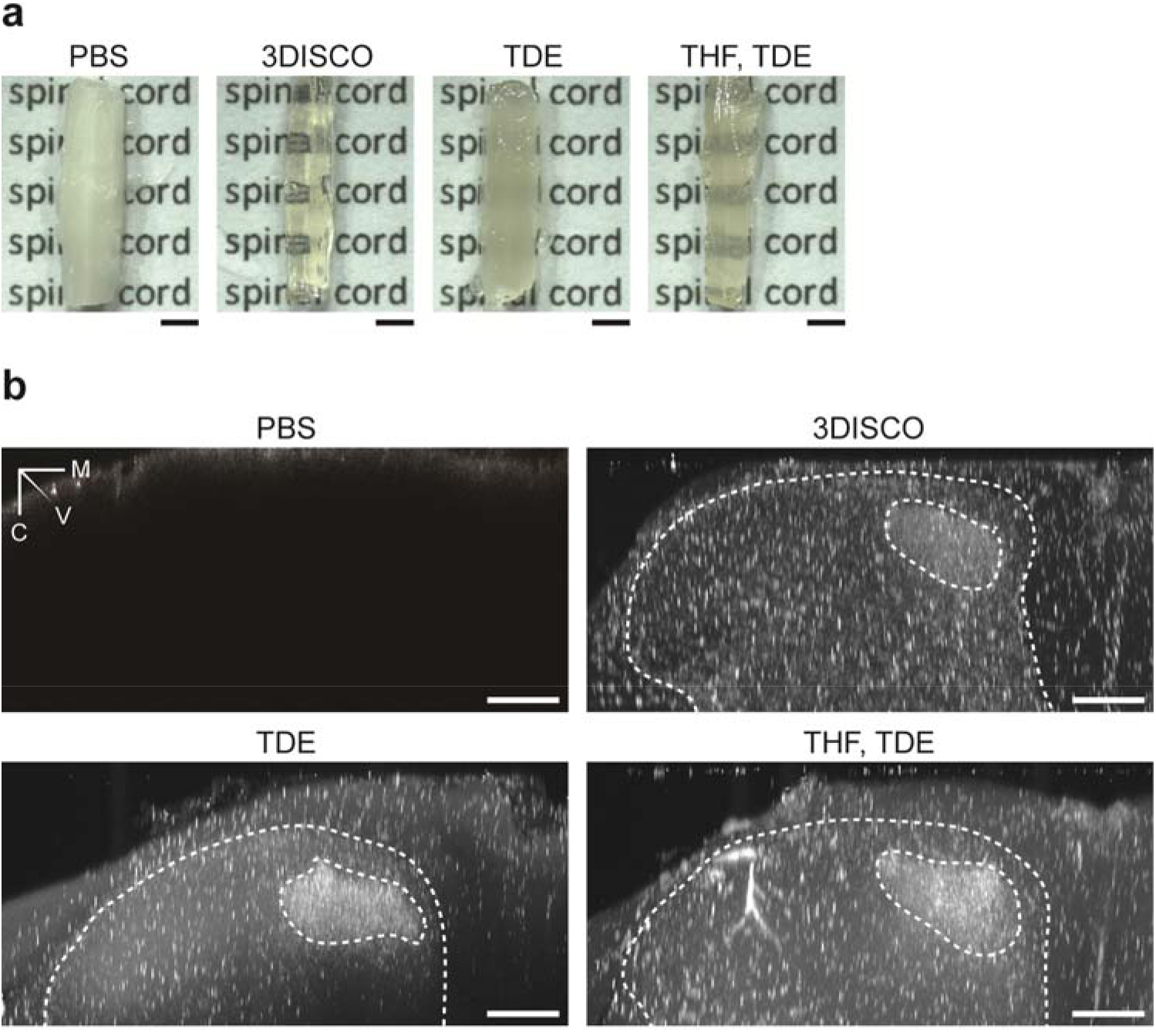
Volumetric imaging of CTB-labelled afferent terminals in the spinal cord dorsal horn. **(a)** Stereoscopic images of spinal cords immersed in PBS or optically cleared using 3DISCO, THF/TDE, or TDE alone. Scale bars: 1 mm. **(b)** Transverse digital slices of spinal cord dorsal horns immersed in PBS or optically cleared using 3DISCO, THF/TDE, or TDE alone. Dashed lines mark the dorsal grey/white matter border. CTB-labelled terminal fields of fibres innervating the left second hind paw digit are encircled. Scale bars: 100 μm. Abbreviations: PBS, phosphate buffered saline; TDE, 2’2 thiodiethanol; M, medial; V, ventral; C, caudal.

**Figure 5-2.**
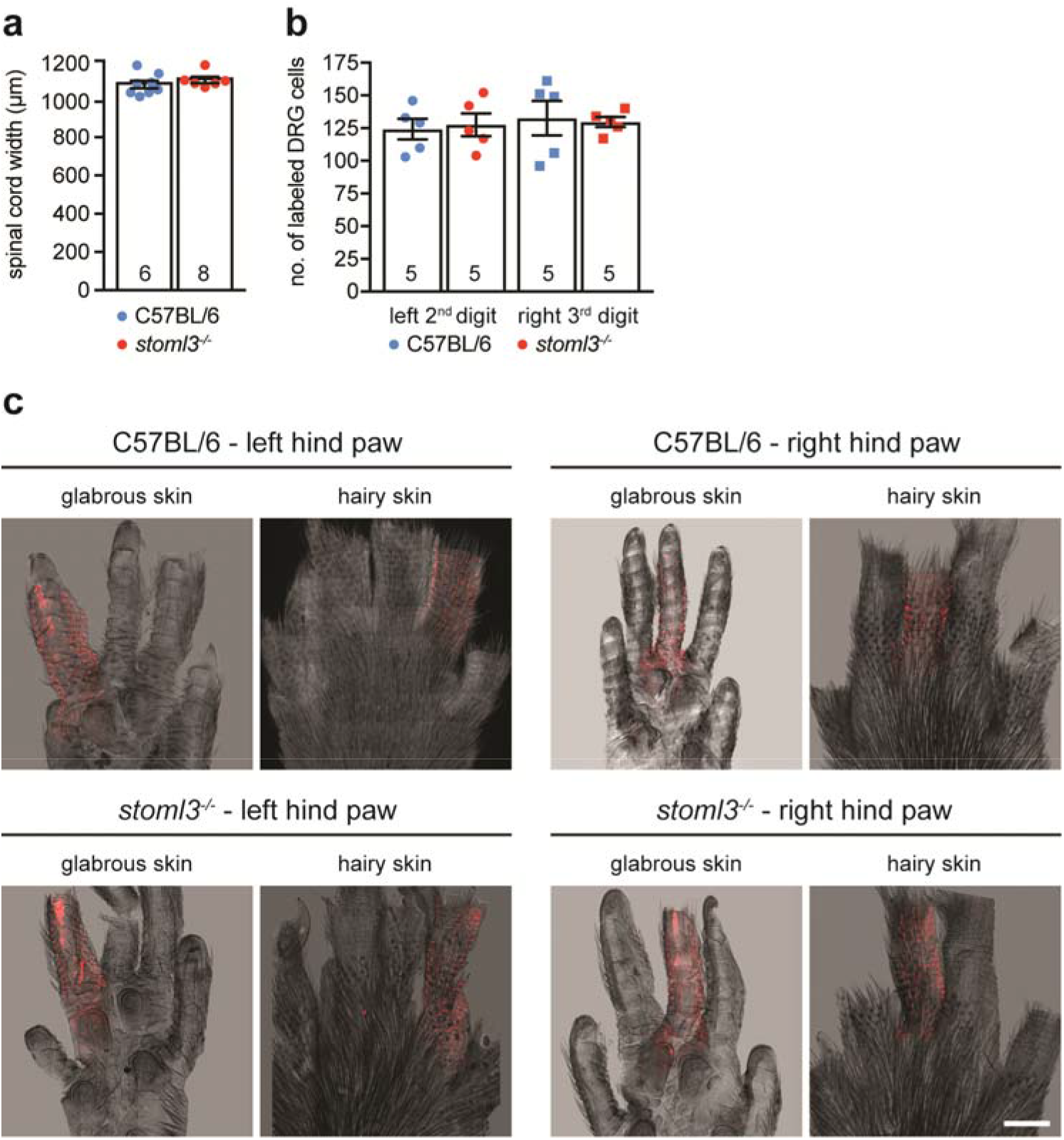
Control experiments ensuring reliable CTB injection performance. **(a)** Width of the spinal cord dorsal horn in *stoml3* mutant (red, n = 6) and control (blue, n = 8) mice. Individual data points and mean values ± SEM are shown. Data was analysed using a two-tailed unpaired t-test. **(b)** Number of CTB-labelled neurons in left (circles) and right (squares) DRGs 3 to 5 in *stoml3* mutant mice (red, n = 5) as compared to control (blue, n = 5) mice. Individual data points and mean values ± SEM are shown. Data was analysed using a two-tailed unpaired t-test. **(c)** Representative images of CTB-labelled skin of the left and right hind in *stoml3* mutant and control mice after subcutaneous injection. Scale bars: 1.5 mm. Abbreviation: DRG, dorsal root ganglion.

**Figure 5-3.**
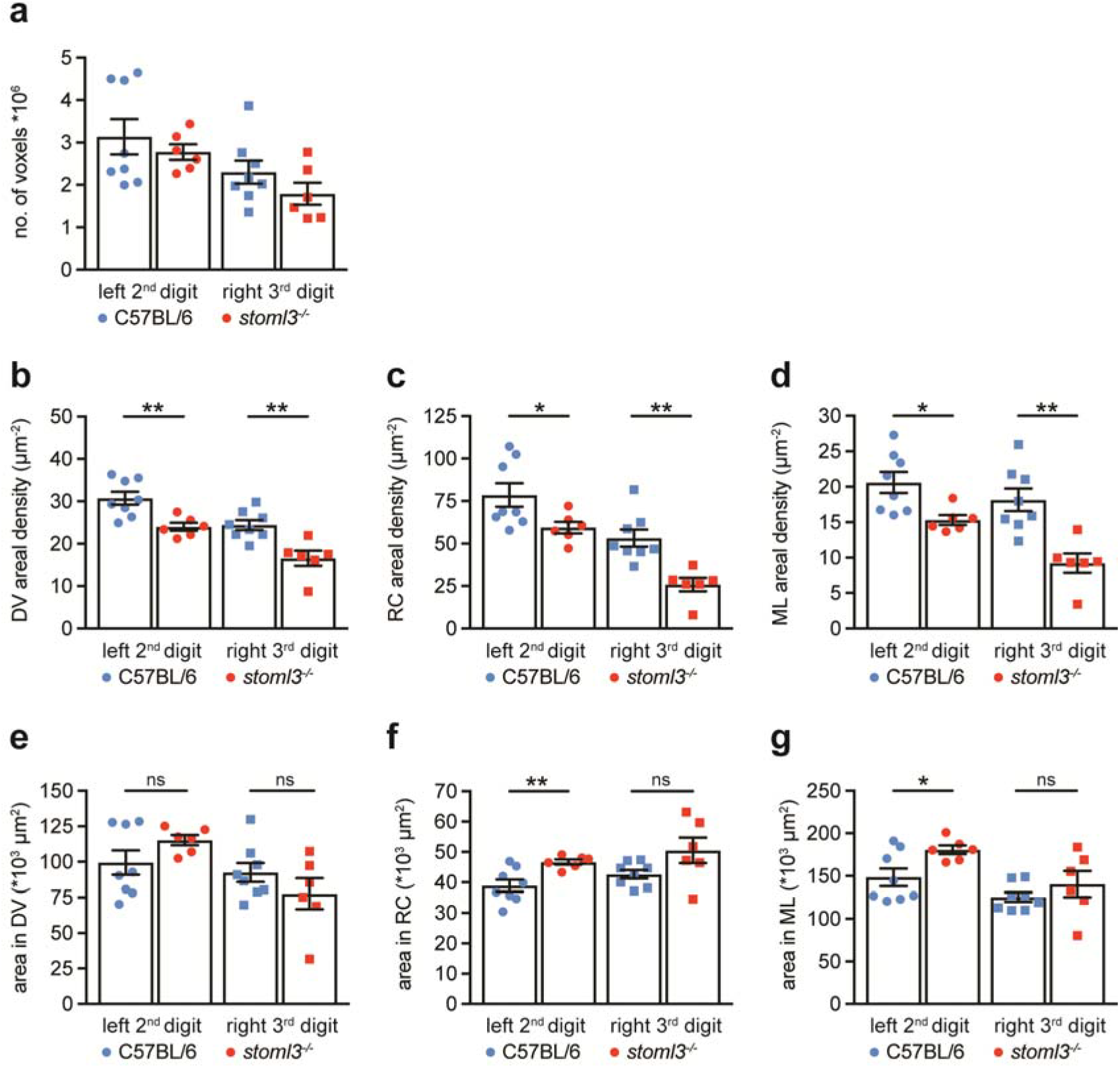
Density measurements of spinal terminal fields in *stoml3* mutant and control mice. **(a)** Numbers of voxels representing CTB-labelled terminals of fibres innervating the left (circles) and right (squares) hind paw digit, respectively, in *stoml3* mutant (red, n = 6) as compared to control (blue, n = 8) mice in tiled image stacks. **(b,c,d)** Areal densities of spinal terminal fields of fibres innervating the left (circles) and right (squares) hind paw digit, respectively, in *stoml3* mutant (red, n = 6) as compared to control (blue, n = 8) mice in **(b)** dorso-ventral, **(c)** rostro-caudal, and **(d)** medio-lateral summed projections. **(e,f,g)** Areas occupied by pixels representing CTB-labelled terminals of fibres innervating the left (circles) and right (squares) hind paw digit, respectively, in *stoml3* mutant (red, n = 6) as compared to control (blue, n = 8) mice in **(e)** dorso-ventral, **(f)** rostro-caudal, and **(g)** medio-lateral summed projections. Individual data points and mean values ± SEM are shown. Data sets were compared using two-tailed unpaired ‘t-tests. Abbreviations: DV, dorso-ventral; RC, rostro-caudal; ML, medio-lateral.

**Figure 6-1.**
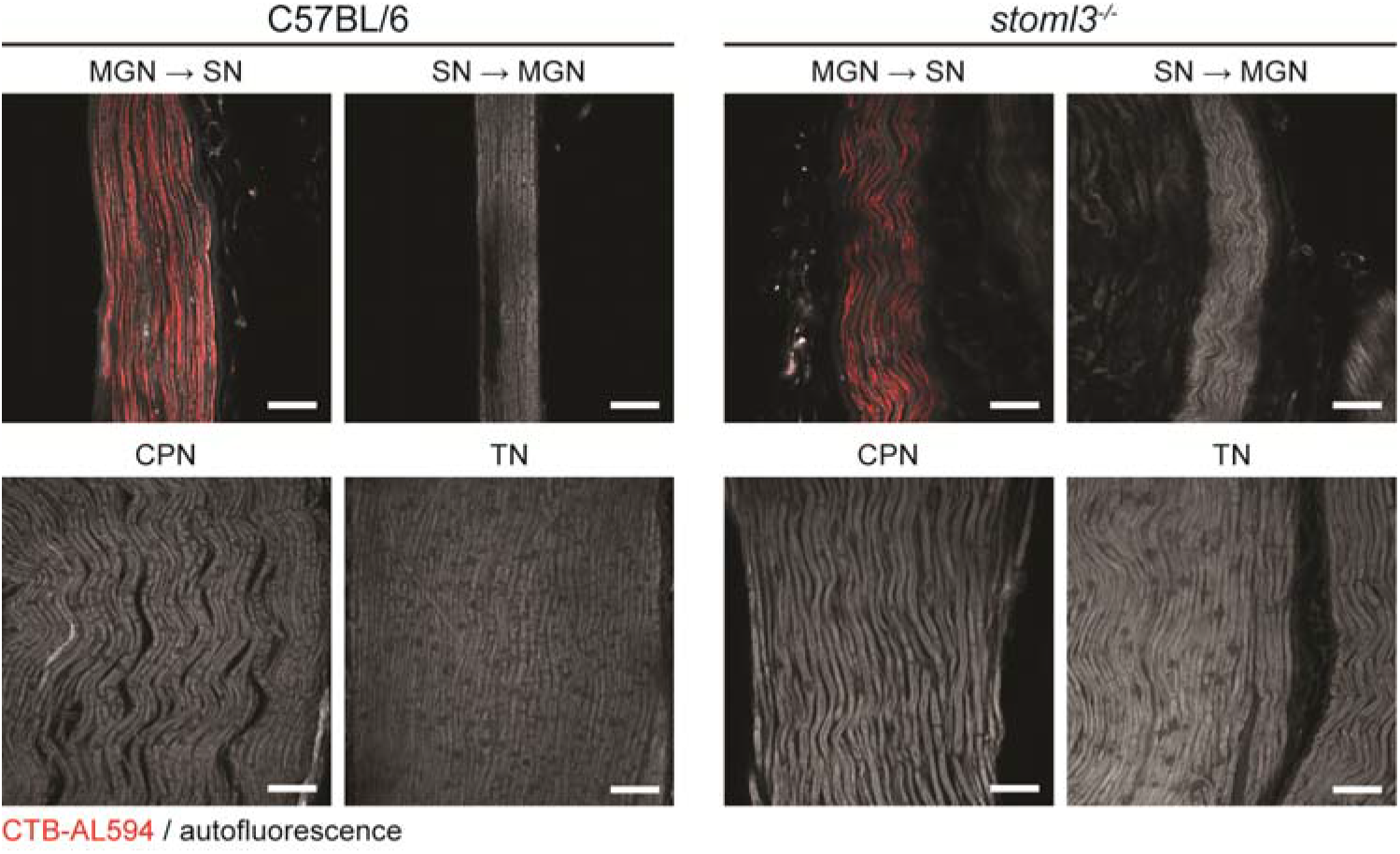
Peripheral nerves after intraneural CTB-injection into the cross-anastomosed gastrocnemius nerve innervating the skin. CTB-labelled myelinated muscle afferents (red) redirected towards the skin (MGN → SN) after intraneural tracer injections distal to the cross-anastomosis site in *stoml3* mutant and control mice. Myelinated sural nerve afferents redirected towards the gastrocnemius muscle muscle (SN → MGN) as well as CPN and TN afferents are not labelled in both control and *stoml3* mutant mice. Autofluorescence is shown in grey. Scale bars: 50 μm. Abbreviations: MGN → SN, muscle afferents from the medial gastrocnemius nerve redirected towards the skin; SN → MGN, cutaneous afferents from the sural nerve redirected towards the gastrocnemius muscle; CPN, common peroneal nerve; TN, tibial nerve.

